# A genome-wide relay of signalling-responsive enhancers drives hematopoietic specification

**DOI:** 10.1101/2022.03.22.485307

**Authors:** B. Edginton-White, A Maytum, S.G. Kellaway, D.K. Goode, P. Keane, I. Pagnuco, S.A. Assi, L. Ames, M Clarke, P.N. Cockerill, B. Göttgens, J.B. Cazier, C. Bonifer

## Abstract

Developmental control of gene expression critically depends on distal cis-regulatory elements including enhancers which interact with promoters to activate gene expression. To date no global experiments have been conducted that identify their cell type and cell stage-specific transcription stimulatory activity within one developmental pathway and in a chromatin context. Here, we describe a high-throughput method that identifies thousands of differentially active cis-elements able to stimulate a minimal promoter at five stages of hematopoietic progenitor development from embryonic stem cells, which can be adapted to any ES cell derived cell type. Exploring this new resource, we show that blood cell-specific gene expression is controlled by the concerted action of thousands of differentiation stage-specific sets of cis-elements which respond to cytokine signals that terminate at signalling responsive transcription factors. Our work presents a major advance in our understanding of developmental gene expression control in the hematopoietic system and beyond.

---

The blueprint for the developmental regulation of gene expression is encoded in our genome in the form of cis-regulatory elements that exist as nuclease hypersensitive sites in chromatin. These elements are scattered over large distances and integrate multiple intrinsic and extrinsic signals regulating the activity of transcription factors (TFs) that bind to such elements^1, 2^. TFs and TF encoding genes together with their targets form gene regulatory networks (GRNs) that define the identity of a cell. TFs together with chromatin remodellers/modifiers form multi-molecular complexes that assemble on cis-regulatory elements and interact with each other within intranuclear space to activate gene expression^2^. To answer the question of how one GRN transits into another in development, it is essential (i) to identify and characterize the full complement of cell type and cell stage-specific transcription regulatory elements, (ii) to identify the TFs binding to them and (iii) to understand how they respond to external cues.

For several decades, reporter gene assays have been used to define cis-regulatory elements as either enhancers and promoters, with enhancers being able to increase transcription from promoters independent of their orientation^3,4^. However, assessing enhancer activity within a chromatin context is more difficult. Studies inserting individual enhancer-promoter combinations at different genomic locations revealed that the local chromatin environment strongly influences gene expression, a phenomenon named a genomic position effect. Such effects can typically only be overcome if the full complement of cis-regulatory elements is present on a transgene, making the analysis of individual elements difficult as their deletion make transgenes again susceptible to position effects^5^. The deletion of individual elements within a gene locus can uncover enhancer function, but often misses developmental stage-specific elements, at least in part due to functional redundancy with neighbouring elements. Therefore, multiple surrogate markers have been identified that correlate with a high activity of the gene linked to the respective cis-regulatory element, including DNaseI hypersensitivity, TF binding, histone acetylation/mono-methylation and enhancer transcription^3,6–11^. None of these features, alone or in combination was fully predictive of enhancer activity with TF binding being the best predictor^12,13^. Consequently, functional assays remain essential to ascertain whether any given element can stimulate transcription in a chromatin context.

During embryonic development, definitive blood cells including hematopoietic stem cells develop from mesoderm derived endothelial cells within the dorsal aorta^14,15^. The specification of hematopoietic cells in the embryo and hematopoietic cell differentiation in the adult have served as important model to reveal general principles of the control of gene expression in mammalian development^16^. The most important TFs together with signals such as cytokines controlling different developmental stages are known, and most intermediate cell types have been identified. Moreover, in vitro differentiation of human and mouse embryonic stem cells (ESCs) recapitulates embryonic hematopoietic development, thus facilitating deep molecular analysis into the developmentally-controlled transition of gene regulatory networks (GRNs)^17^.

We previously reported a multi-omics analysis revealing dynamic GRNs that are specific for each of the major stages of blood cell specification. We identified the locations and dynamic activities of sets of cis-regulatory elements associated with developmental gene regulation and based on this data, and uncovered important pathways, such as Hippo signalling that are required for blood cell formation^18–20^. However, major questions are still open. Whilst we could correlate chromatin alterations with dynamic gene expression^21^, our data did not provide functional evidence for which cis-regulatory elements have enhancer activity, how they are controlled at different developmental stages, how extrinsic signals control their activity and importantly, which TFs mediate signalling responsiveness.

In the work presented here, we developed a novel high-throughput method identifying thousands of enhancer and promoter elements specifically active at defined stages of blood cell specification in a chromatin environment and correlate their activity with gene expression in the same cells. We used mouse ESCs to be able to integrate our results with our previously global multi-omics collected data. However, the method can be expanded to any cell type that can be differentiated from ESCs and can easily be adapted to human ESCs. We show that the same elements exist as active chromatin in vivo. Finally, we identified cytokine responsive enhancer elements, and for one cytokine, VEGF, characterized the TFs mediating its activity within the GRN driving blood cell development. Our work provides an important resource for studies of hematopoietic specification and highlights the mechanisms of how extrinsic signals program a cell type-specific chromatin landscape driving hematopoieetic differentiation.

## Results

### Establishing a high-throughput method for functional enhancer testing in a chromatin environment

We identified functional enhancer elements in the chromatin of mouse embryonic stem cells and their differentiated progeny representing different stages of hematopoietic specification (Figure 1a)^18,19^. The first stage analysed here consists of FLK1-expressing hemangioblast-like cells^22^ which have the ability to differentiate into cardiac, endothelial and hematopoietic cells and which are purified from embryoid bodies (EBs) after day 3 of culture. These cells are then placed into blast culture and form a mixture of (i) hemogenic endothelium 1 (HE1) which expresses a low level of RUNX1, (ii) hemogenic endothelium 2 (HE2) cells which up-regulate RUNX1 and CD41 but are still adherent and (iii) blast-like hematopoietic progenitor cells (HP) that underwent the endothelial-hematopoietic transition (EHT). Using the differential expression of specific surface markers (KIT, TIE-2 and CD41) each cell type can be purified to near homogeneity.

**Figure 1:**
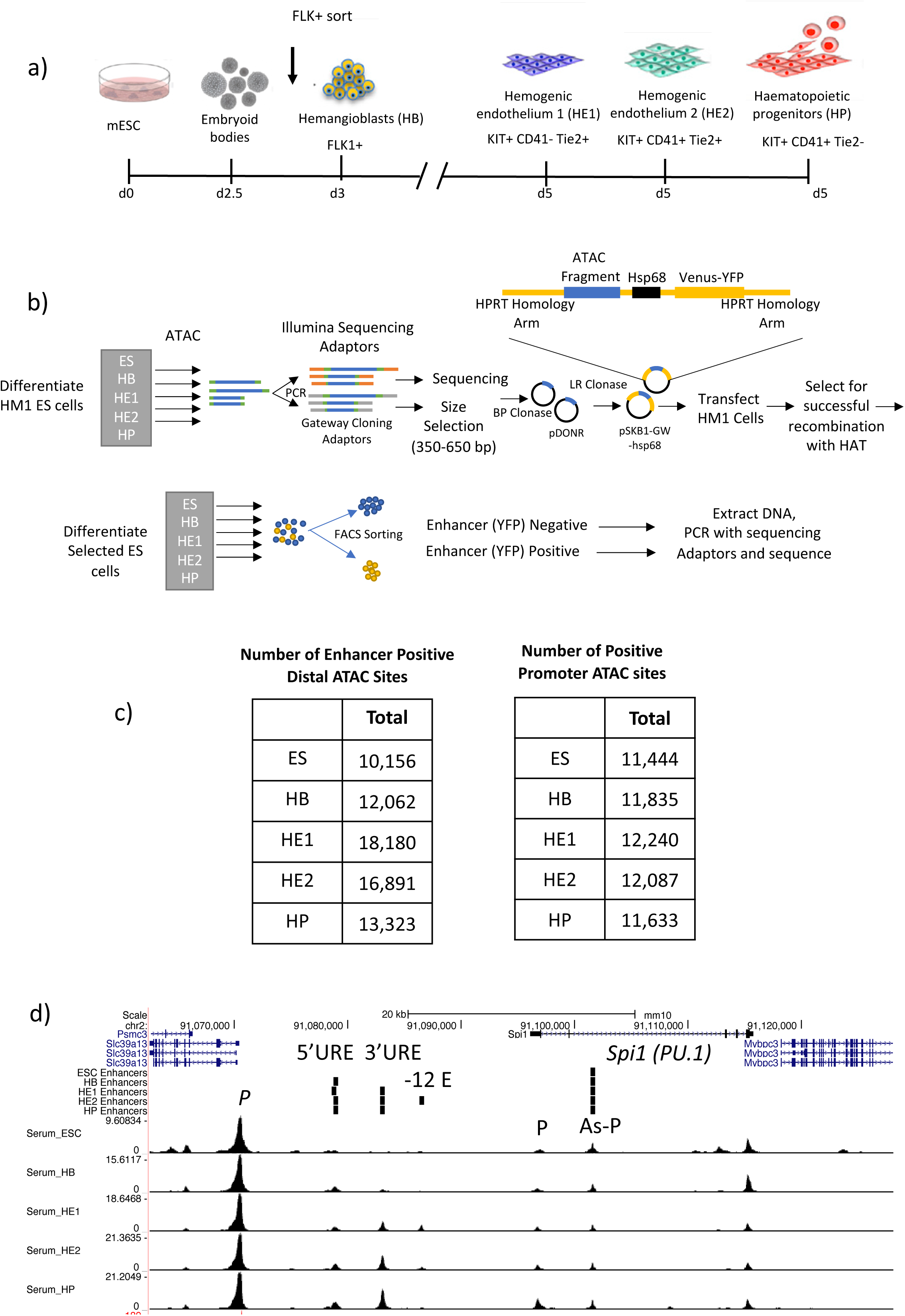
Establishing a high-throughput method for functional enhancer testing in a chromatin environment. a) Depiction of the ES cell differentiation system and the cell types analysed. b) Overview over the screening procedure using ATAC-Seq fragments from 5 different cell stages. c) Total number of distal and promoter ATAC-sites scoring positive in two replicates of the assay. d) an example of well characterized enhancers found in the *SPI1* (PU.1) locus ^24^ found in our screen showing enhancer screen and ATAC-Seq data.

Figures 1 and S1 show the enhancer identification pipeline which is based on the system developed by Wilkinson et al., 2013^23^ and was adapted for high-throughput. Essentially, we differentiate cells, sort them into different developmental stages as shown in Figure 1a, purify ATAC-Seq fragments for each differentiation stage, clone them into a targeting vector to generate a fragment library which is then integrated into a defined target site in the *HPRT* locus carrying a minimal promoter to drive a reporter gene (Venus-YFP) (Figure 1b). We then differentiate cells and purify cells from each stage of development for reporter activity measurements (Figure S1 a,b). Cell populations expressing high, medium and low YFP levels are isolated together with YFP negative cells (Figure S1b). Fragment inserts are sequenced after amplification using barcoded primers recognising the ATAC linkers. Sequences then undergo a rigorous filtering against different criteria as detailed in Figure S1c. Our screen was conducted in two replicates identifying several hundred-thousand fragments with transcription-stimulatory activity. 22% - 31% of fragments were located within annotated distal elements (Figure S1d), covering more than 70,000 enhancer-positive ATAC sites across all stages (Figure 1c left panel, Figure S1e). The remaining fragments were promoter sequences which were classified as being 1.5 kb within an annotated transcription start site (Figure 1c, right pane, Figure S1d). An example for enhancer annotation is shown in Figure 1d, depicting the well-characterized *Spi1* (PU.1) locus, which captures all previously identified enhancer elements together with their known stage-specific activity^24^. To our surprise, most ATAC-fragments displaying stimulatory activity (see scheme in Figure S2a, Figure S1d, left panel) overlapped with ATAC-sites containing negatively scoring fragments, indicating that the vast majority of captured open chromatin regions can regulate transcription.

We next integrated our enhancer data with previously published chromatin immunoprecipitation (ChIP)-Seq data characterizing histone modifications at cis-regulatory elements in the same experimental system^18^. About 30% to 60% of all enhancer sites overlap with H3K27Ac regions (Figure 2a, S2b). We find a significant overlap of our positive but not our negative/unknown enhancer ATAC sites with the VISTA enhancer database which describes 1061 functionally identified enhancers^25^ (Figure 2b, S2c). The size distribution of positive and negative (non-scoring) fragments was the same (Figure S2d) indicating no size selection for active fragments. We also show that the most enhancer positive ATAC fragments overlap with open chromatin sites found in purified hemogenic endothelium and endothelial cells from day 9.5 and day 13.5 mouse embryos^26,27^ indicating that the same elements are active in vivo (Figure S2e). Finally, the comparison with previously collected TF ChIP data^18,20,28,29,30^ shows that between 17% (in HB) and 76% (in HP cells) of all enhancer fragments are bound by ubiquitous and differentiation stage-specific TFs, depending on the number of available ChIP experiments for each stage (data not shown).

**Figure 2:**
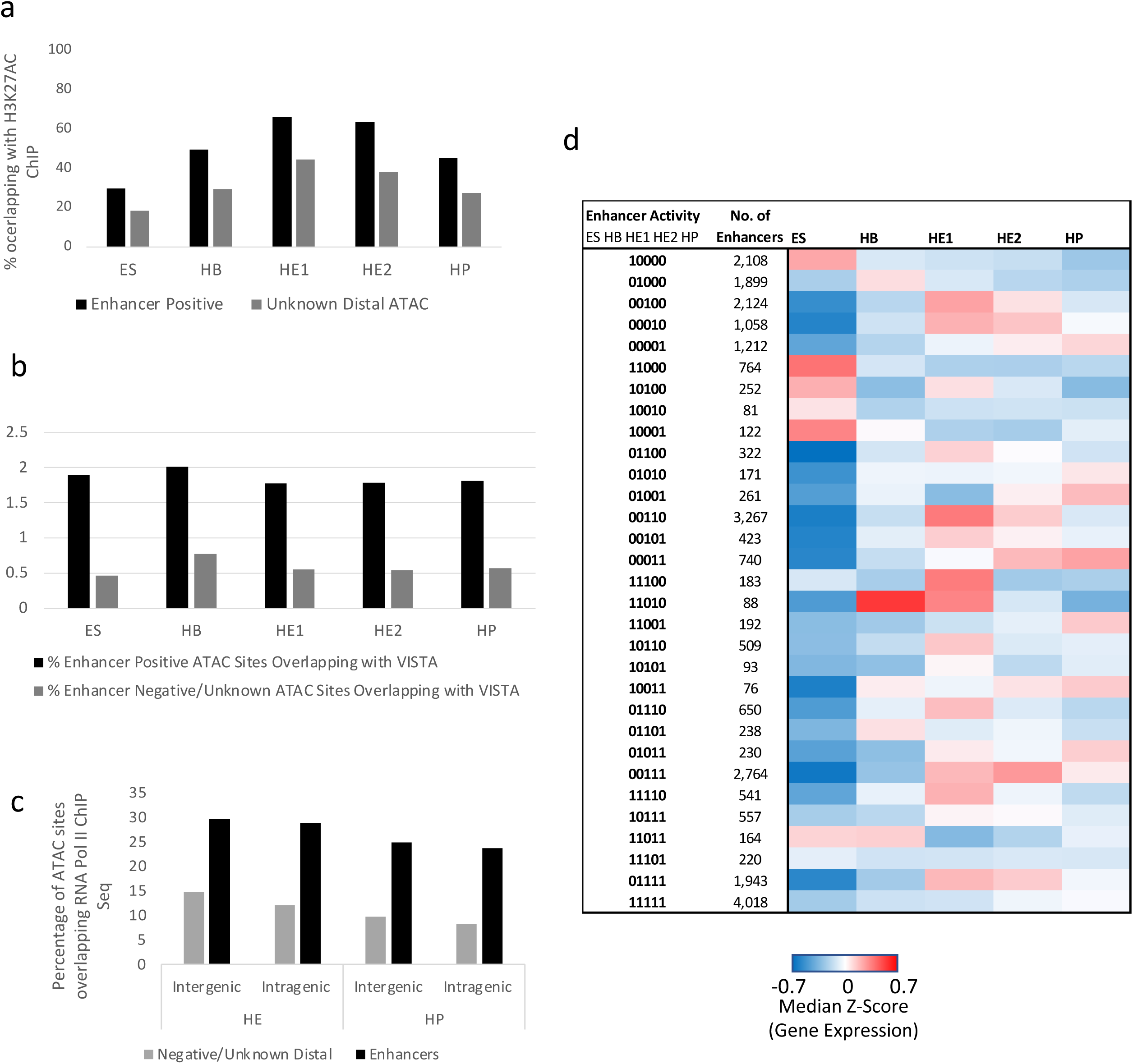
Characterization of enhancer features and association of cell stage specific enhancer activity with cell stage-specific gene expression. a) Upper panel: Percentage of functionally identified enhancer elements overlapping with histone H3 lysine 27 (H3K27Ac) peaks as compared to inactive elements or where activity is not known. Data from ^18^. b) Overlap of newly discovered enhancer elements with known enhancers from the VISTA database ^25^. Percentage of functionally identified intergenic and intragenic enhancer elements overlapping with RNA polymerase II binding sites in the hemogenic endothelium and HP cells. Data from^29^. d) Presence or absence of enhancer activity at the different developmental stages expressed as binary code (0 = inactive; 1 = active) and sorted by the activity pattern across differentiation. The heatmap depicts the activity of the genes associated with these elements. Enhancer - promote association was determined by using the union of HiC data from ES and HPC7 (HP) cells together with co-regulation data determined by ^21^ in total 21671 elements. All promoters not covered by these data (5599) were associated by being the nearest to the enhancer element.

It was reported that active enhancers are bound by RNA-Polymerase II (Pol II) and are transcribed^8,31^. This feature formed the basis for high-throughput transient transfection assays (STARR-Seq)^32^ which identified thousands of elements active in this assay in one cell type. However, it is unclear how many of these elements can function in a chromatin environment and how their cell stage-specific activity is regulated. To examine the correlation between the ability of distal elements identified in our study to drive RNA production during development we examined whether they were capable of binding Pol II using a previously published data-set from ESC generated HE and HP cells^29^. Figure 2c shows that up to 30% of identified enhancers are associated with PolII binding.

### Sequential stages of hematopoietic specification are defined by distinct enhancer sets

The genomic information for tissue-specific gene expression manifests itself in the activity pattern of distal regulatory elements^18,33^. An important feature of our method is therefore the identification of developmental stage-specific enhancer activity. To assign genes to regulatory elements, we used publicly available HiC and co-regulation data^21,28,34^ and examined how gene expression correlated with enhancer activity by generating a binarized matrix cataloguing enhancers as active (1) and inactive (0) at each of the five differentiation stages (Figure 2d). This analysis shows that (i) HE1 and HE2 are highly related cell types, (ii) enhancer activity during development is largely continuous and (iii) stage-specific gene expression is strongly associated with stage specific enhancer activity.

We then determined for each differentiation stage, which TF motifs and motif combinations were associated with enhancer activity (Figure 3 and S3). We first examined the motif content of distal ATAC-Seq sites at specific differentiation stages^18^. In line with previously published ChIP data^18,20,28^, motif patterns are associated with the binding of stage-specific TFs (Figure S3a), with those for hematopoietic TFs such as RUNX1 or PU.1 being enriched in hematopoietic progenitor cells and those for the HIPPO signalling mediator TEAD and SOX factors enriched in hemogenic endothelium. This pattern was also seen with stage-specific activity of enhancers but not with promoter fragments (Figure S3c). In contrast, ubiquitously active enhancer fragments displayed a similar motif signature at all developmental stages, reinforcing the notion that tissue-specificity is encoded in distal elements. The motif for the Zn++ finger factor CTCF was enriched in active distal enhancer but not promoter fragments. It was recently shown in differentiating erythroid cells that dynamically bound CTCF cooperates with lineage-specific TFs bound to distal elements to interact with promoters^35^. Our data are consistent with this finding.

**Figure 3:**
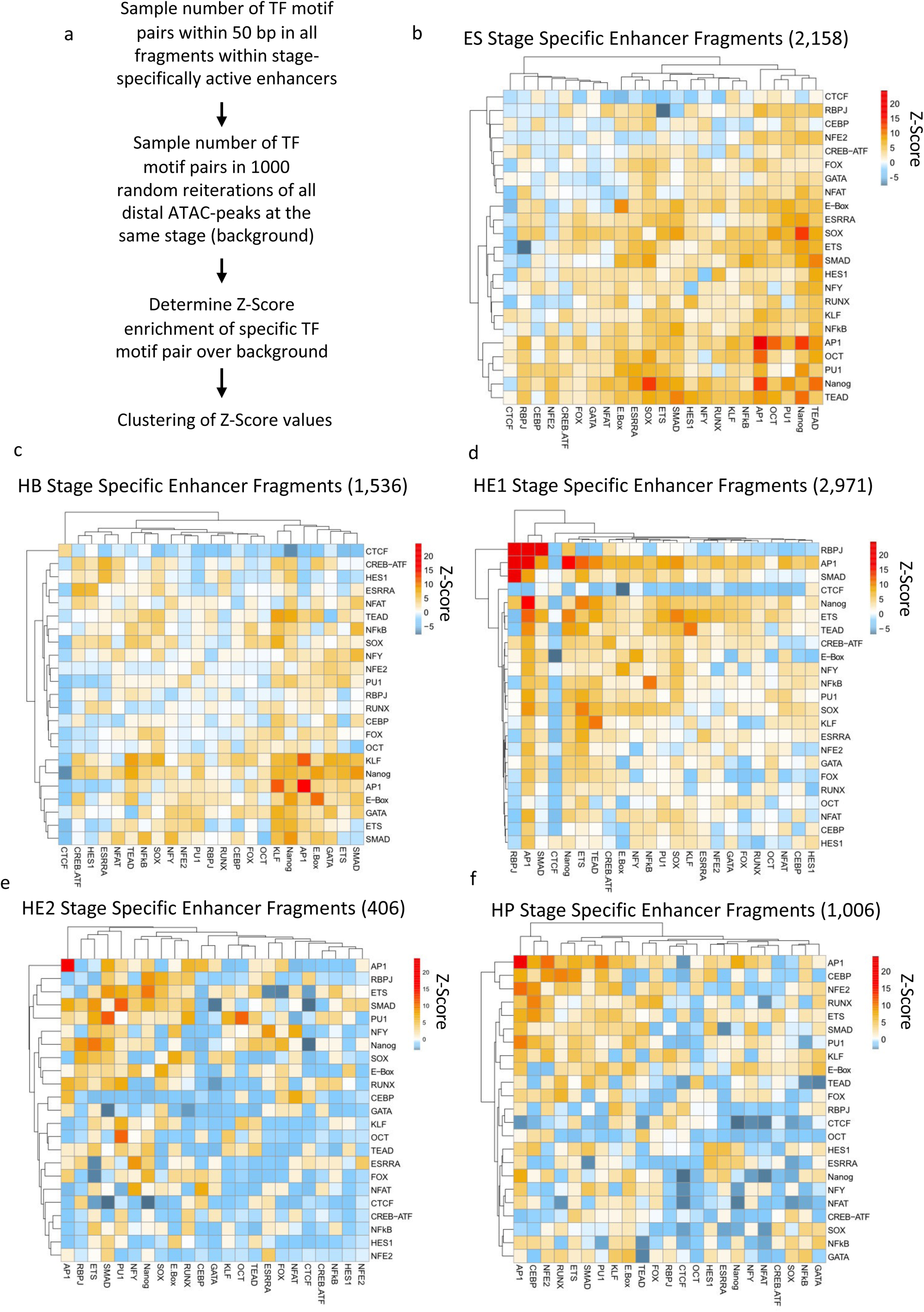
Stage-specific enhancer elements show a specific TF motif co-localization pattern. a) Overview over motif co-localization analysis pipeline, a - e): Heatmaps depicting Z-Score enrichments for five cell differentiation stages as indicated, together with the number of fragments analyzed. Binding motifs for the indicated TFs are listed on the right and the bottom of the heat-map.

We next asked whether stage-specific enhancer activity was correlated with a pattern of cooperating TFs. To this end, for each developmental stage we performed a motif co-localization analysis which examines whether specific binding motif pairs located within 50 bp of each other were enriched in stage-specific enhancer fragments as compared to all open chromatin sites (Figure 3a). Motifs with a 100% overlap were removed. Stage-specific enhancer activity correlated with enriched colocalizations of motifs for developmental-stage specific TFs. In line with their generally open chromatin structure^36^, ESC-specific enhancers showed a great variety of paired motifs with those for the pluripotency factors NANOG/SOX2/OCT4 showing co-localization. Interestingly, AP-1 motifs showed a very high co-occurrence both with itself and other motifs, potentially linking their activity to signalling processes (Figure 3b). AP-1 homo-typic motif associations were also found at HB-specific enhancer fragments but co-localization with pluripotency factor motifs was lost (Figure 3c). The HE1 stage showed a strong enrichment in RBPJ, AP-1 and SMAD motif pairs, with AP-1 motifs colocalizing with high frequency with most other factors (Figure 3d). The number of HE2-specific enhancers was low as it is a transitory stage. Here, PU.1 motifs show increased co-localization with SMAD and OCT motifs (Figure 3e). As expected from previous ChIP studies^37^, we find a co-localization of motifs for hematopoietic TFs such as C/EBP, RUNX, PU.1 and GATA at the HP-stage. AP-1 motifs again co-localized with a variety of other motifs, including C/EBP, RUNX1 and PU1 (Figure 3f). Although generally enriched in enhancer fragments at all stages (Figure S3c), CTCF motifs were not significantly paired with any other motif.

### Identification of cytokine-responsive enhancer elements

Cell differentiation involves extracellular signals which alter growth and differentiation state by changing gene expression. Signalling molecules such as cytokine receptors, integrins and kinase molecules are well characterized, but less is known of how different signals are integrated at the level of the genome. We have limited information about which cis-regulatory elements can respond to signals, which TF combinations are involved and how they cooperate to ensure that the genome responds to outside signals in a coordinated and balanced fashion. Our global cis-element collection allows us to answer these pivotal questions.

To this end, we employed a serum-free in vitro differentiation system^38^ that is based on the sequential addition of growth factors such as BMP4, VEGF and hematopoietic cytokines. We tested how individual cytokines affected the differentiation profile and the open chromatin landscape of HE1, HE2 and HP cells sorted as described in Figure 4a. The comparison of the chromatin signature of cells differentiated in serum and under serum-free conditions (Figure 4a), showed that around 80% of cis-elements seen in cells from serum-free culture overlapped in both conditions, demonstrating the reproducibility of our differentiation system. However, for HP cells we noticed changes in the bulk open chromatin landscape which affected the distal elements and not the promoters. This result indicates that although the cellular identity seemed to be largely preserved in sorted cells expressing the right combination of surface markers, the difference in the signalling environment exerted a strong effect on the chromatin landscape.

**Figure 4.**
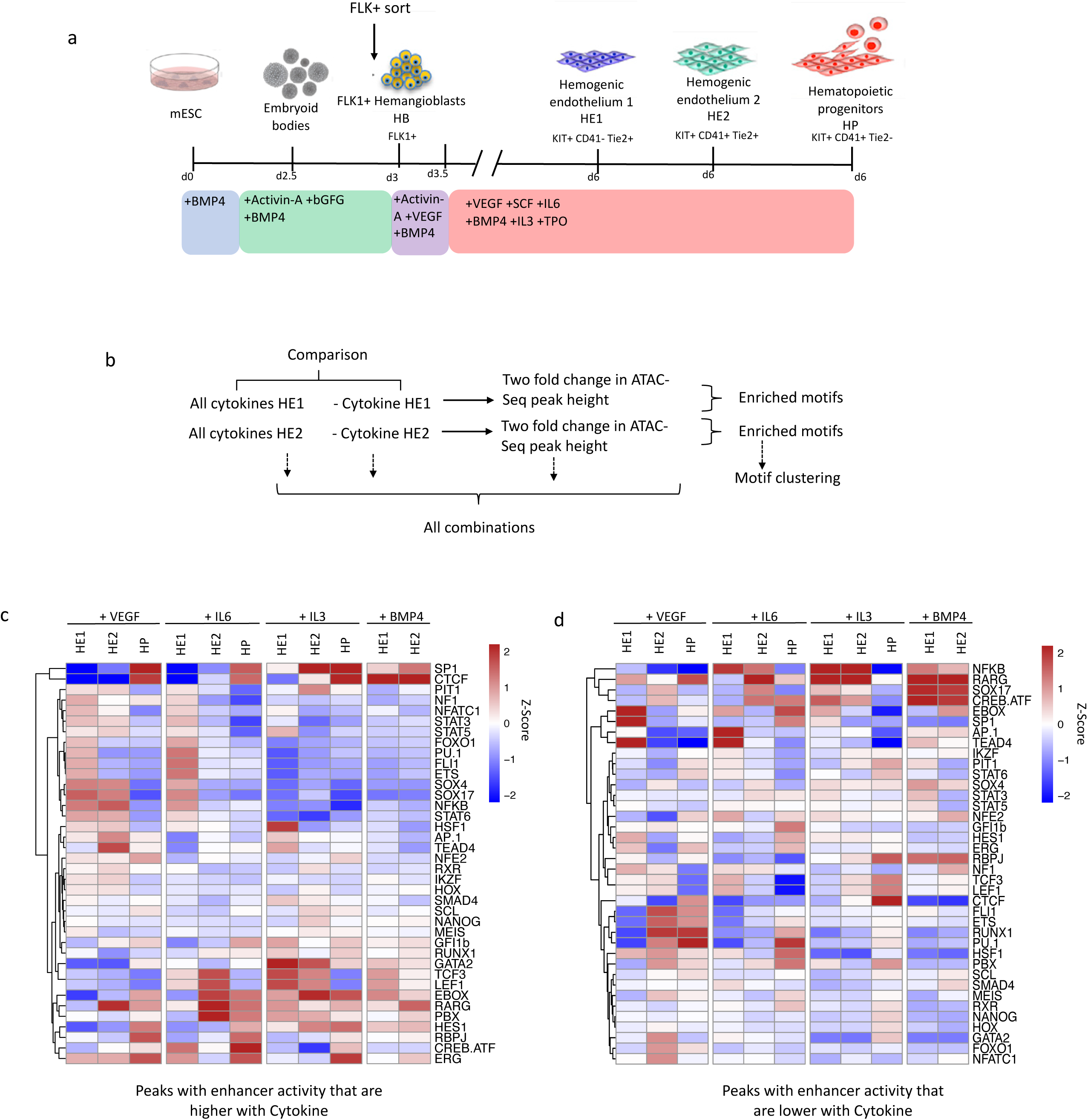
Identification of cytokine-responsive enhancer elements, a) Overview of serum-free in vitro blood progenitor differentiation modified from ^38^. Specific cytokines were withdrawn of left out at the beginning of blast culture b) Overview of motif enrichment analysis strategy, c,d) : TF binding motif enrichment in distal ATAC peaks harbouring enhancer activity that are increased (c) or decreased (d) in the presence of the indicated cytokines.

To elucidate the role of the different cytokines on chromatin programming, we differentiated cells in the presence and absence of specific cytokines (in this case BMP4, VEGF, IL-6 and IL-3), sorted the different cell types and examined which open chromatin regions changed at least two-fold in response to cytokine withdrawal. In total, more than 10000 unique open chromatin regions were up or down regulated (Figure S4b) which included both distal elements and promoters. We next examined, which TF binding motifs were enriched in cytokine-responsive ATAC-Seq peaks harbouring enhancer activity (Figure 4b - d). Alteration of cytokine conditions had a profound influence on chromatin programming. The absence of BMP was incompatible with HP formation and open chromatin regions with enhancer activity containing SMAD, HOX, RAR and NOTCH motif signatures were lost in HE. This finding is in keeping with these factors being required to form the HE. We noticed that the presence of VEGF led to a loss of peaks with a hematopoietic motif signature in HE2/HP, such as RUNX1, FLI1, GATA and PU.1 motifs. In contrast to VEGF, the presence or absence of BMP4, IL-6 and IL-3 did not influence the proportion of generated HE1, HE2 and HP cells as compared to the all-cytokine condition (Figure 5a). A similar motif enrichment pattern was found when all open chromatin regions were analysed (Figure S4d,e).

**Figure 5.**
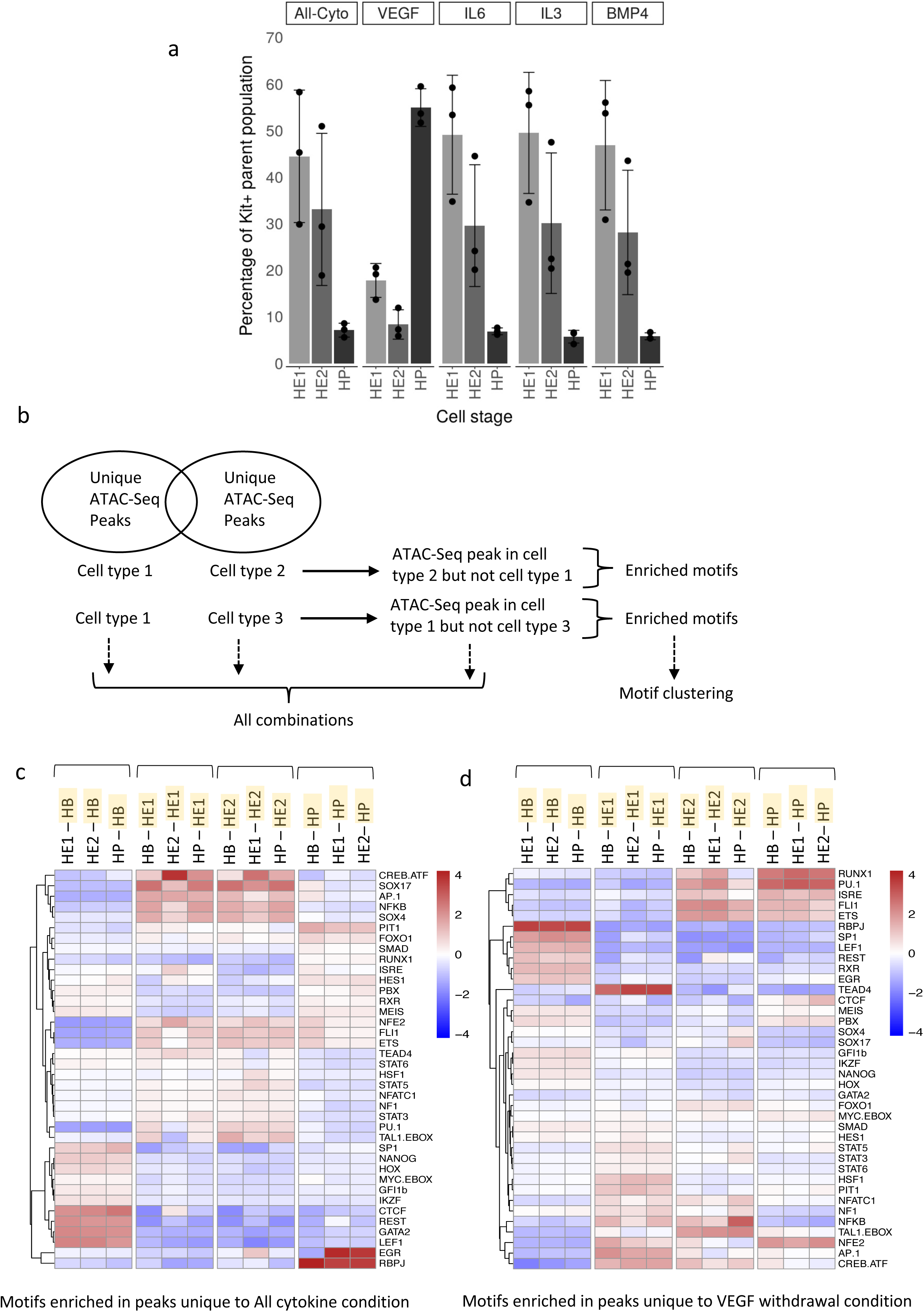
The presence of VEGF suppresses multipotent progenitor (HP) development. a) Proportion of HE1/HE2/HP cells within the differentiation culture in the absence of the indicated cytokines as measured by FACS. Error bars represent the standard deviation from n=3 with the spread being indicated by dots. b-d) Un-supervised motif clustering analysis examining enrichment of the TF binding motifs in all ATAC peaks. b) motif enrichment strategy; c) motifs enriched in peaks of specific cell types (highlighted in yellow) unique to All Cytokine Condition, d) motifs enriched in peaks of specific cell types (highlighted in yellow) unique to the VEGF withdrawal condition. TF binding motifs are depicted at the right of the panel.

### The balance between hematopoietic and endothelial development is regulated by VEGF-responsive cis-regulatory elements

VEGF had the greatest influence on the formation of hematopoietic progenitor cells (Figure 5a). We therefore examined its role in regulating enhancer activity in more detail. VEGF withdrawal cultures contained an altogether smaller number of cells but an increased proportion of HP cells (Figure S5a). Moreover, VEGF removal at different time points of blast culture did not influence the proportion of HE1 cells but led to an inverse correlation of HE2 and HP cell numbers (Figure S5b), indicating that this cytokine impacted on the EHT. These results are consistent with previous observations showing that the receptor for VEGF, FLK1, is essential for the formation of blood islands as they originate from hemogenic endothelium cells^39^ but once hematopoietic cells are formed, cells become dependent on hematopoietic cytokines. However, the molecular basis of this finding, i.e. which genomic events are responsible for this phenomenon has so far been unclear.

Our ATAC-Seq analysis found 6860 chromatin regions carrying enhancer elements that responded to VEGF. To identify VEGF responsive TFs, we conducted a supervised motif clustering analysis that highlighted cell type specific motif enrichments (Figure 5a–c). This analysis indicates that the withdrawal of VEGF activates enhancers with a hematopoietic motif signature with RUNX1 and PU.1 motifs. HP cells in VEGF cultures maintain an enrichment of motifs for RPBj and HES1 which are mediators of NOTCH signalling^40^. Moreover, in contrast to +VEGF cultures, ATAC peaks in −VEGF HE1 cells were strongly enriched in TEAD motifs together with binding motifs for factors linked to inflammatory signalling (AP-1, NFkB and CREB/ATF) which has been shown to be important for stem cell development^41^. A similar motif signature in VEGF-withdrawal cultures was also seen when ATAC sites were directly compared (Figure S5).

### The VEGF response involves the TEAD – AP-1 axis

We next examined which specific TFs were connected to endothelium-specific gene expression and VEGF responsiveness. Distal ATAC-Seq peaks specific for HE cells derived from cultures containing VEGF were enriched for SOX, E-BOX, AP-1 and TEAD TF motifs, indicating a distinct endothelial TF motif signature. This was also true for ATAC peaks with enhancer activity in the HE, pointing to a role of the HIPPO signalling mediator TEAD and the MAP kinase signalling mediator AP-1 in regulating enhancer activity. We previously identified HIPPO signalling as being crucial for HE development^18^ and others have shown that shear stress activating the TEAD partner YAP via RHO-GTP induces the formation of HSCs from the HE via activation of *RunxX1*^42^. Our ChIP data show that AP-1 and TEAD indeed bind to such motifs in the HE^20^ (Figure 6c) but once HP cells have formed, TEAD binding is lost^18^. To examine the role of these TFs in more detail, we identified enhancer elements which were active specifically in the HE and were bound by these factors. The *Galnt1* enhancer fitted this criterion (Figure 6a, c). We created several ES reporter cell lines carrying the wild-type sequence and TF binding motif mutations (Figure 6b).

**Figure 6.**
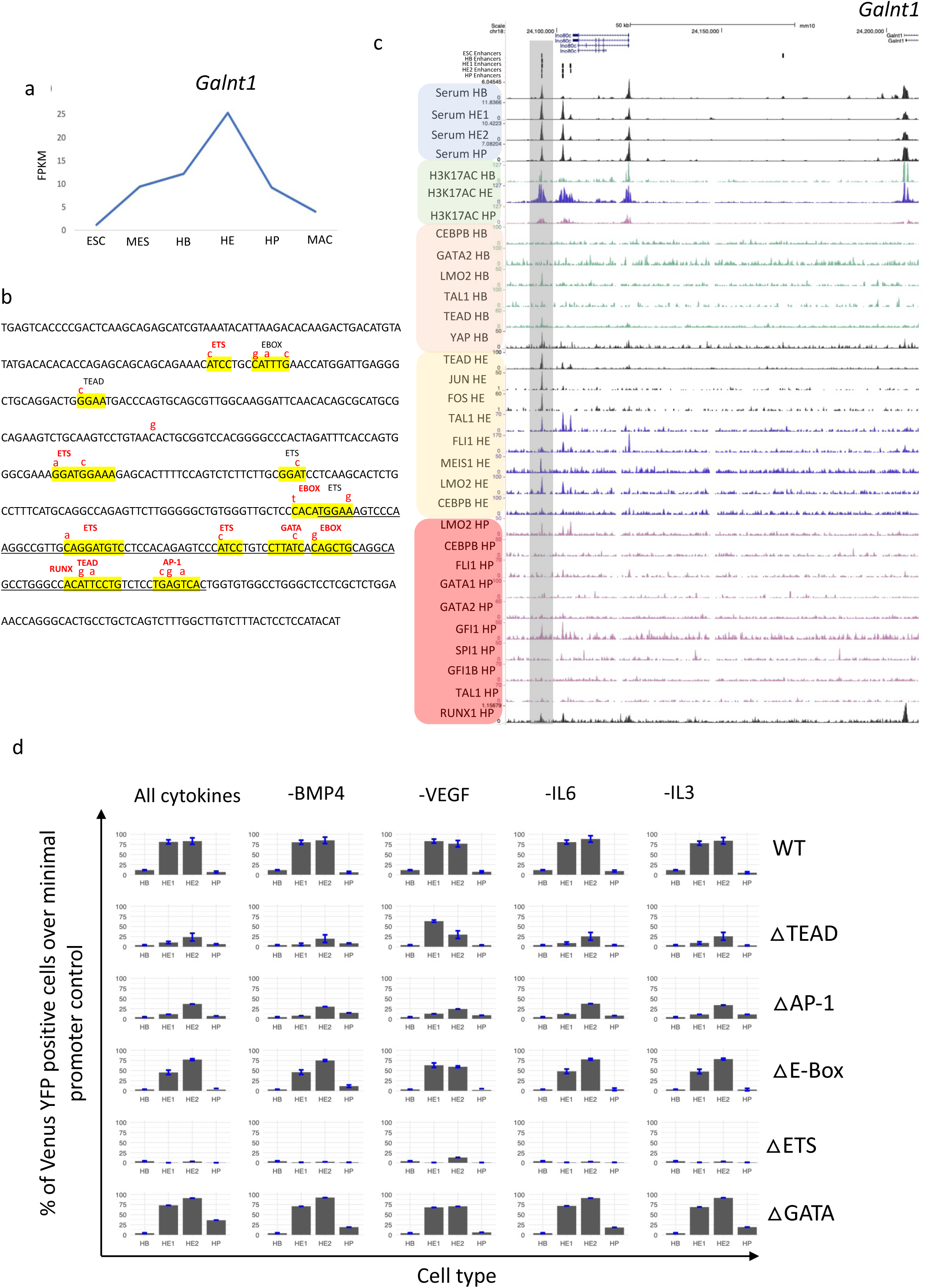
HE-specific expression of the *Galnt1* gene is mediated by an upstream enhancer element binding TEAD and AP-1. a) HE-specific expression of *Galnt1* (data from ^18^. b) Sequence of the *Galnt1* enhancer with TF binding motif mutations indicated in red. Strong consensus sequences are highlighted in red. c) UCSC genome browser screenshot showing ChIP data from Goode et al. ^18^ and Obier et al. ^20^ and enhancer activity measured in this study. The *Galnt1* enhancer and its promoter are highlighted with a grey bar. The *Ino80* gene is constitutively expressed (data not shown) and *Galnt1* was shown to be co-regulated with the presence and absence of a DHS at this enhancer ^21^. Note that this enhancer is bound by RUNX1 in the HP which binds to a site overlapping with the TEAD::AP-1 element. d) Reporter gene activity driven by wild type and mutated *Galnt1* enhancer elements in the presence and absence of the indicated cytokines. Error bars represent the standard deviation from n=3.

We previously reported that TEAD and AP-1 often show a specific spacing between binding sites where binding is interdependent^20^. The expression of a dominant negative (dn) FOS peptide blocking all AP-1 DNA binding reduced the binding of TEAD at such sites, suggesting that HIPPO and MAP kinase signalling interface there. The *Galnt1* enhancer carries one of these composite AP-1/TEAD motifs (Figure 6b). In addition, dnFOS induction reduced *Galnt1* mRNA expression specifically in the HE (Figure 6b), indicating that AP-1 activates the element and is required for the binding of TEAD factors within this composite module. The analysis of *Galnt1* gene expression in the presence and absence of VEGF showed that the cytokine causes a delay in down-regulation of this gene which is consistent with the delayed formation of HP cells (Figure S6c).

To examine the *Galnt1* enhancer response to cytokines at four differentiation stages (HB – HP), we first measured the activity of the intact enhancer element (Figure S6c) as compared to the promoter control by using flow cytometry to assay the number of YFP positive cells. The activity profile of the enhancer mirrored the gene expression profile in the presence of all cytokines (Figure 6d). We also measured median YFP florescence (Figure S6d). The analysis of mutant enhancer elements revealed that the binding motifs for TEAD, ETS and AP-1 were the most important for enhancer activity in the HE whereas the mutation of others had no impact (Figures 6d, S6d and data not shown).

The absence of BMP4, IL-6 and IL-3 had little effect on the number of YFP+ cells but in the absence of VEGF median reporter activity across all cells was reduced in the HE (Figure S6d). Moreover, the mutation of the TEAD binding sites led to an increase in reporter activity in HE1 cells, suggesting that here TEAD restricts enhancer activity. Taken together, these data demonstrate that VEGF modulated *Galnt1* enhancer activity is regulated by the balanced interplay of AP-1 and TEAD TFs which integrate two signalling pathways.

### VEGF blocks the upregulation of *RUNX1* at the chromatin and gene expression level via up-regulating *Notch1/Sox17*

The data shown above demonstrate that VEGF interferes with the EHT and progenitor formation. Both processes are crucially dependent on the TF RUNX1^19^ which is required to activate hematopoietic genes and to repress endothelial genes in cooperation with the TF GFI1^43^. Analysis of the TFs binding to the *Galnt1* enhancer revealed that in HP cells it is bound by RUNX1 and GFI1 (Figure 6c)^18^ at a site overlapping the TEAD motif but TEAD no longer binds there, suggesting a factor exchange and the establishment of repressive RUNX1 complex. To examine why the presence of VEGF leads to *Galnt1* upregulation, we examined the chromatin structure of *Runx1* in the presence and absence of VEGF (Figure 7a). In the presence of VEGF multiple distal DHSs of *Runx1* fail to form and 5 of these elements score in our enhancer assay, including the previously characterized +23 kb enhancer (2)^44^. Of note, several, of these elements, in particular enhancer 3, are bound by TEAD and AP-1 as well.

**Figure 7.**
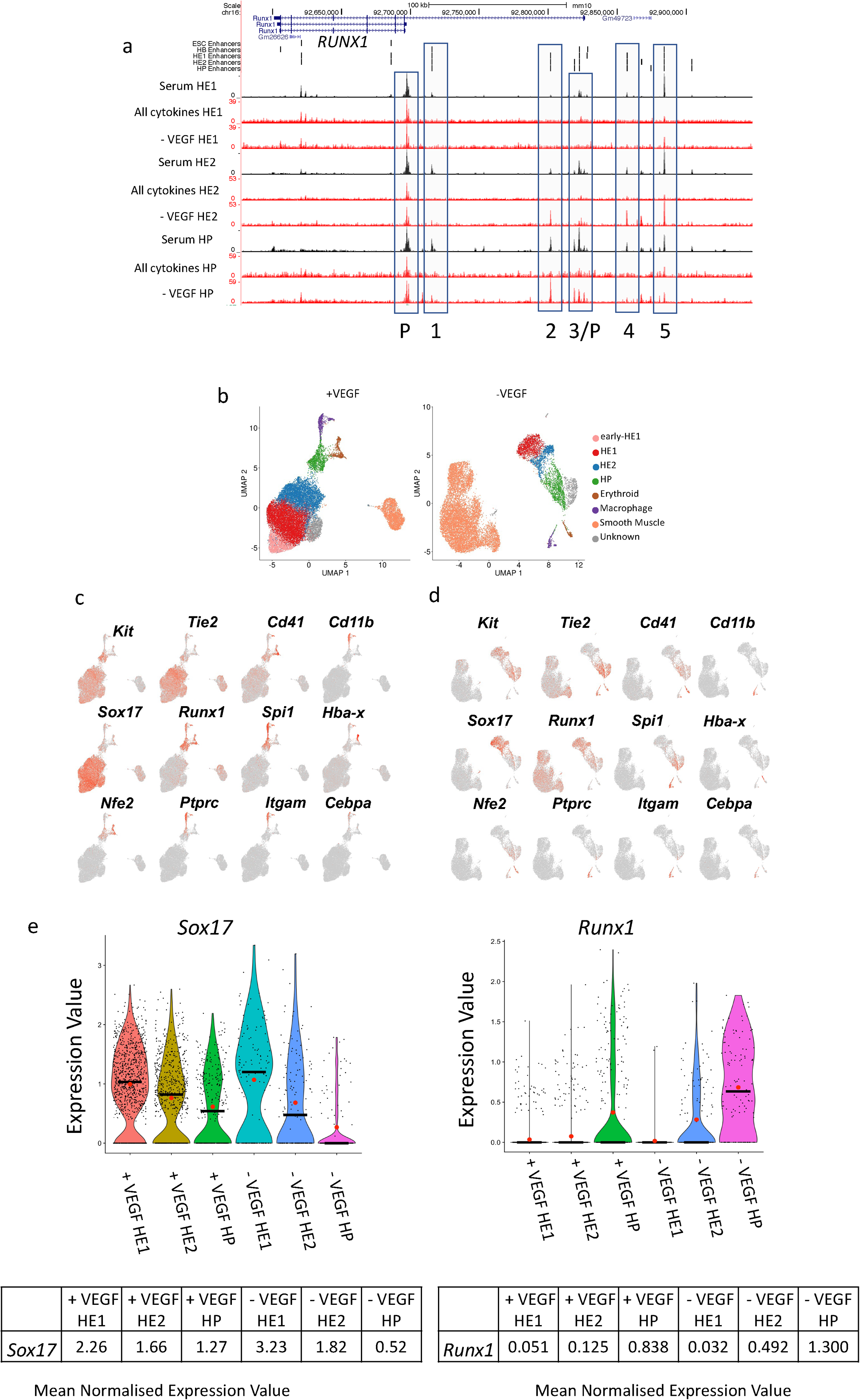
VEGF withdrawal shifts the proportion of endothelial / smooth muscle and hemogenic endothelium / blood progenitor cells by blocking *Runx1* enhancer activity. a) UCSC genome browser screenshot depicting open chromatin regions and enhancer locations in the *Runx1* locus under the indicated culture conditions. Enhancer (numbered 1 - 5) and promoter elements are indicated by a blue box. b) Single cell gene expression analysis with purified HE1 and HE2 populations in the presence and absence of VEGF. UMAP clustering analysis was performed after all data were pooled, c-d) gene expression levels projected on the clusters shown in b). e) Expression levels of *Sox17* and *Runx1* in the indicated cell types in the presence and absence of VEGF calculated from single cell expression data.

The results described so far suggested that the reduced ability of VEGF containing culture to undergo the EHT was a result of a failure of *Runx1* enhancer activation and a concomitant failure of its transcriptional upregulation in HE2. Although we profiled chromatin in FACS purified cells, these alterations could still be caused by shifts in cell composition. We therefore studied VEGF-mediated changes in gene expression in HE1/2 cells at the single cell level (Figure 7b-f, S7a – e). Without VEGF the overall numbers of HE1/HE2/HP cells were reduced (Figure 7b) but the population showed an increased proportion of HP cells, together with an increase in the proportion of smooth muscle cells (Figure 7b). However, the cellular identity and the overall differentiation trajectory were not altered (Figure S7e). This shift was in concordance with the lack of down-regulation of endothelial genes such as *Sox17* and *Tie2* (Figure 7 c,d,e) and a lack of upregulation of *Runx1* in HE2 and HP cells (Figure 7e, right panel). The balance between endothelial and hematopoietic gene expression is controlled by the TF SOX17 which represses *Runx1*^45,46^. After the EHT, RUNX1 together with GFI1 binds to *Sox17* and *Notch1* enhancers ^18,43,47,48^ thus forming a feed forward loop that drives EHT progression. *Sox17* and *Runx1* are thus expressed in a mutually exclusive fashion with the former being high before the EHT^46^ and then being downregulated and the latter being upregulated from a low level during the EHT^47^. This balance shifted after VEGF was withdrawn (Figure 7 d-f). We therefore conclude that VEGF signalling interferes with the activation of *Runx1* enhancers in the HE, driving gene expression required for the EHT.

To investigate whether VEGF-responsive enhancers were connected to VEGF regulated genes, we paired each element that required VEGF activation in HE1, HE2 and HP with its respective gene as shown in Figure 2d. We then integrated this data with their expression as measured in our single-cell experiments (Figure S7f). This analysis uncovered that at high-stringency analysis, between 10% and 19% of VEGF responsive enhancers are linked to VEGF responsive up- or down-regulated genes, thus directly linking alteration in chromatin with changes in gene expression. Moreover, the analysis of gene ontology (GO) terms of VEGF dependent genes highlighted angiogenesis as the top pathway. VEGF-responsive down-regulated genes linked to down-regulated enhancer elements include hematopoietic regulator genes such as *Runx1, Klf2, Jun* and *Elf1*, again validating our general approach.

We next examined how VEGF impacted on other signalling factors involved in regulating HP development. The formation of the arterial HE as a source of HP cells is dependent on NOTCH1 signalling^40,49^. VEGF and SOX17 activate *Notch1* and lack of down-regulation of NOTCH1 blocks the EHT^45,46,50,51^. *Notch1* and *Sox17* are down-regulated after the EHT and after upregulation in HP cells, RUNX1 and GFI1 repress *Sox17*^18,43,47,48^, thus forming a feed forward loop driving EHT progression. We examined whether any of the components of this pathway was modulated by VEG. We found that the *Dlk1* gene encoding a repressor of NOTCH1 activity^52^ was strongly upregulated in the absence of VEGF, both at the chromatin and the gene expression level (Figure 8a,b) thus contributing to regulating the switch from an endothelial to a hematopoietic program.

**Figure 8.**
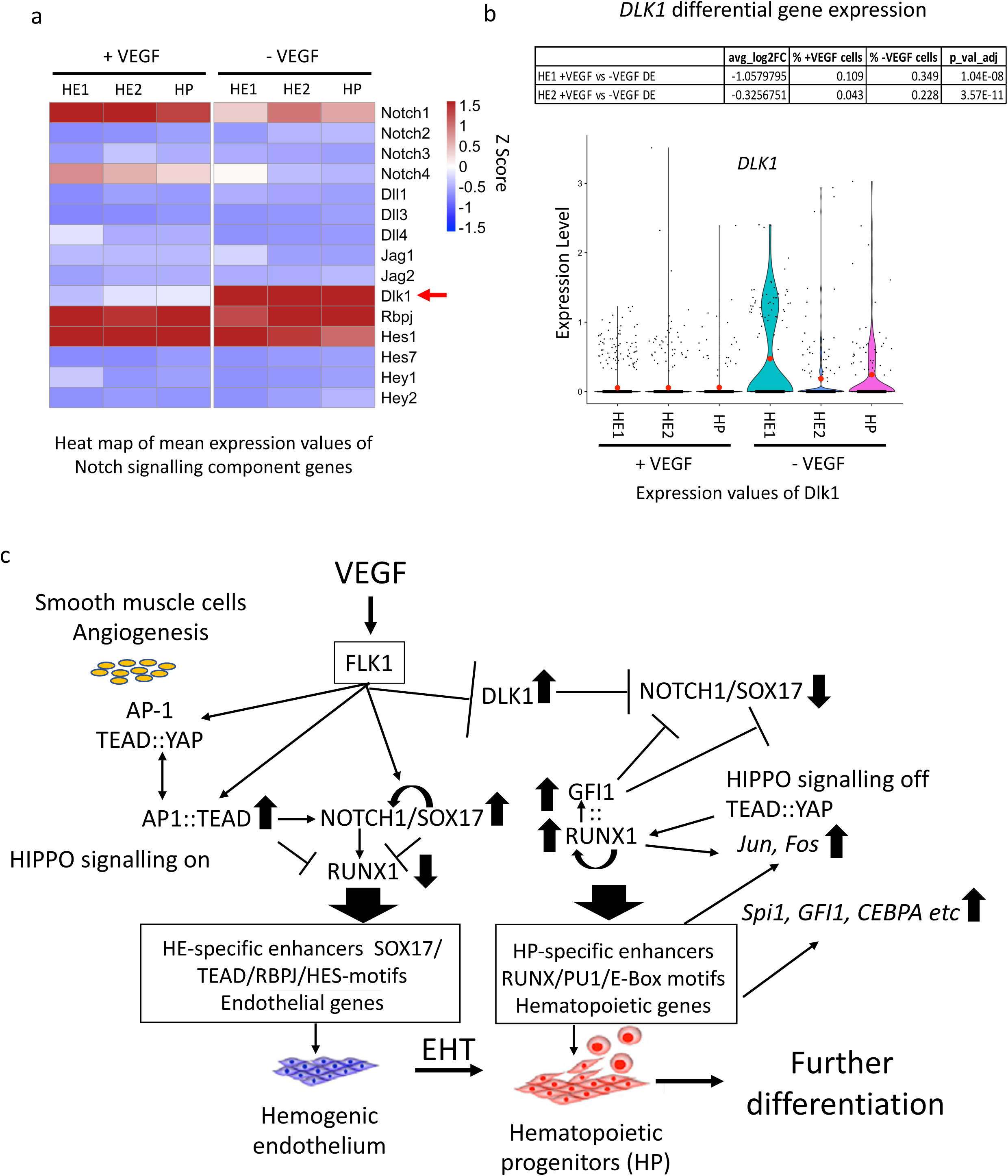
VEGF regulates the balance between endothelial and hemogenic development by controlling NOTCH1. (a) Heatmap showing the expression of NOTCH signalling genes with and without VEGF based on single cell expression data, b) expression of Dlk1 with and without VEGF. c) Core gene regulatory and signalling network regulating blood stem cell emergence. For further details see text.

Taken together, our data show that VEGF signalling in the hemogenic endothelium impacts on a gene regulatory network (Figure 8c) that controls NOTCH1 signalling responsive enhancer elements which bind TEAD and AP-1 and are enriched in NOTCH signature (RBPJ/HES) and SOX motifs. The signalling-responsive activation pattern of these enhancers represents the molecular engine driving the EHT.

## Discussion

Our study introduces a versatile method that identifies developmental stage-specific, functional enhancers for any cell type that can be differentiated from ES cells. Importantly, our method recovers many enhancers described in the VISTA data-base, for example those of the *Spi1* or the *RUNX1* locus^24,44^, but due to its sheer size, the dataset reported here vastly extends such enhancer sets. The enhancer collection described here thus comprises an important resource for gene targeting approaches and for studies of how signals modulate enhancer activity. Once inserted in the targeting site, stage-specifically active single elements can also perform as tracers for different developmental stages of blood cell development. The method can easily be adapted to human cells and examination of conserved sequences showed that more that 20% of such enhancer elements were also found in the human genome (data not shown).

All identified cis-elements tested in our assay are derived from open chromatin and are enriched for motifs of differentiation stage-specific TFs known to be expressed at this stage. Integration with ChIP-Seq data from multiple sources confirms that these factors indeed bind these motifs. The proportion of non-scoring fragments outside of a chromatin region scoring positive was surprisingly small, demonstrating that most open chromatin regions surrounding genes can impact on transcription. Inactive fragments most likely represent enhancer sub-fragments providing information about enhancer sub-structure. Distal elements identified in this study only partially overlap with chromatin features that have been associated with enhancer sequences, such as H3K27Ac. However, enhancers are nucleosome-free and are only flanked by modified histones. Nucleosomes cover 200 bp of DNA, each increasing the sequence space in which a functional element could be located. We therefore believe that assaying the function and sequence composition of open chromatin regions as described here will provide a much more precise tool to home in on those sequences with true transcription-regulatory activity. We see an association of distal elements with sites of RNA-Polymerase binding, confirming that enhancer transcription is wide-spread.

We find that thousands of regulatory elements are connected to extracellular signals and contain binding motifs for signalling responsive TFs. In response to VEGF, thousands of chromatin regions are opened or closed, directly impacting on the expression of their associated genes. Our single cell experiments show that VEGF truly impedes differentiation, with *Runx1* failing to be upregulated, *Notch1* and *Sox17* not being downregulated, the EHT being delayed, and fewer HP cells being formed, thus directly linking signalling dependent enhancer activity to cell fate decisions. Based on this data, we suggest that the activity of most enhancer elements is fine-tuned by signalling. Our data comparing alternate differentiation conditions (serum versus serum-free versus with or without cytokines) show that signalling impact is highly culture dependent, adding another level of complexity which will make it imperative to seek differentiation conditions mimicking those found *in vivo*. In this context it is relevant that AP-1 motifs are enriched in all stage-specific enhancers and co-localize with multiple tissue-specifically expressed TF motifs, with the AP-1:TEAD site in the *Galnt1* enhancer being an example. AP-1 is known to interact with chromatin remodellers to assist in the binding of other TFs^53^, and multiple studies showed that it is involved in cell fate decisions in response to signals. For example, AP-1 binding is essential for signalling dependent chromatin priming during T cell differentiation^54^. Blocking its activity at the HP stage during ES cell differentiation leads to a complete abolition of myelopoiesis^20^. AP-1 therefore represents a ubiquitous axis integrating the genomic response to specific external signals with internal gene expression programs.

In addition to basic insights into enhancer function, our study provided mechanistic insights into how the core gene regulatory and signalling network regulating hematopoietic specification is connected to the genome (Figure 8c). VEGF orchestrates differential enhancer and promoter activity which regulates the balance between the NOTCH1/SOX17 axis establishing HE identity and RUNX1 driving the EHT and hematopoietic development. VEGF signals to AP-1^55,56^, and we previously showed that this complex regulates the balance between hemogenic and vascular smooth muscle, i.e the endothelial fate^20^. AP-1 also interfaces with HIPPO signalling via TEAD TFs. Switching off HIPPO signalling via YAP activation induces *Runx1* expression in response to shear stress^42^ and in endothelial cells promotes angiogenesis and endothelial gene expression^57^. Taken together with these findings, our ChIP, single-cell gene expression and reporter gene studies place the interplay between NOTCH, HIPPO-signalling and AP-1 mediating MAP kinase signalling responsive TFs at specific enhancer elements at the heart of the balance between endothelial or hematopoietic fate. Although most of the network components are known, it was unclear how they interface with the genome and how they are connected within cell specific gene regulatory networks. In this study, we have now identified the factors involved in their regulation and the genomic elements upon which they act, providing a rich resource for studies of identifying the signals required for the activation of the correct gene expression program required for efficient blood cell production.

## Acknowledgements

This research was funded by a project grant from the Biotechnology and Biological Sciences Research Council (BBSRC) to C.B. and J.B. (BB/R014809/1), a BBSRC MiDTP studentship to C.B. for A.M., a BBSRC LoLa grant to C.B and B.G. (BB/I001220/2), as well as grants from Blood Cancer UK (15001) and the Medical Research Council (MR/S021469/1) to CB and P.N.C. We thank Birmingham Genomics for expert sequencing services and the Birmingham Technology Hub for cell sorting facilities.

## Author contributions

B.E-W., A.M., S.K., L.A., D.G. performed experiments and generated as well as analysed data, P.K, S.A.A. and I.P. analysed data, J.B. supervised data analysis and together with P.N.C. helped writing the manuscript, C.B. conceived and directed the study and CB and B.E-W. wrote the manuscript.

## Data availability statement

All genome-wide data as well as code will be available in public repositories when requested / after publication.

## Extended Figure legends

**Extended Figure S1:**
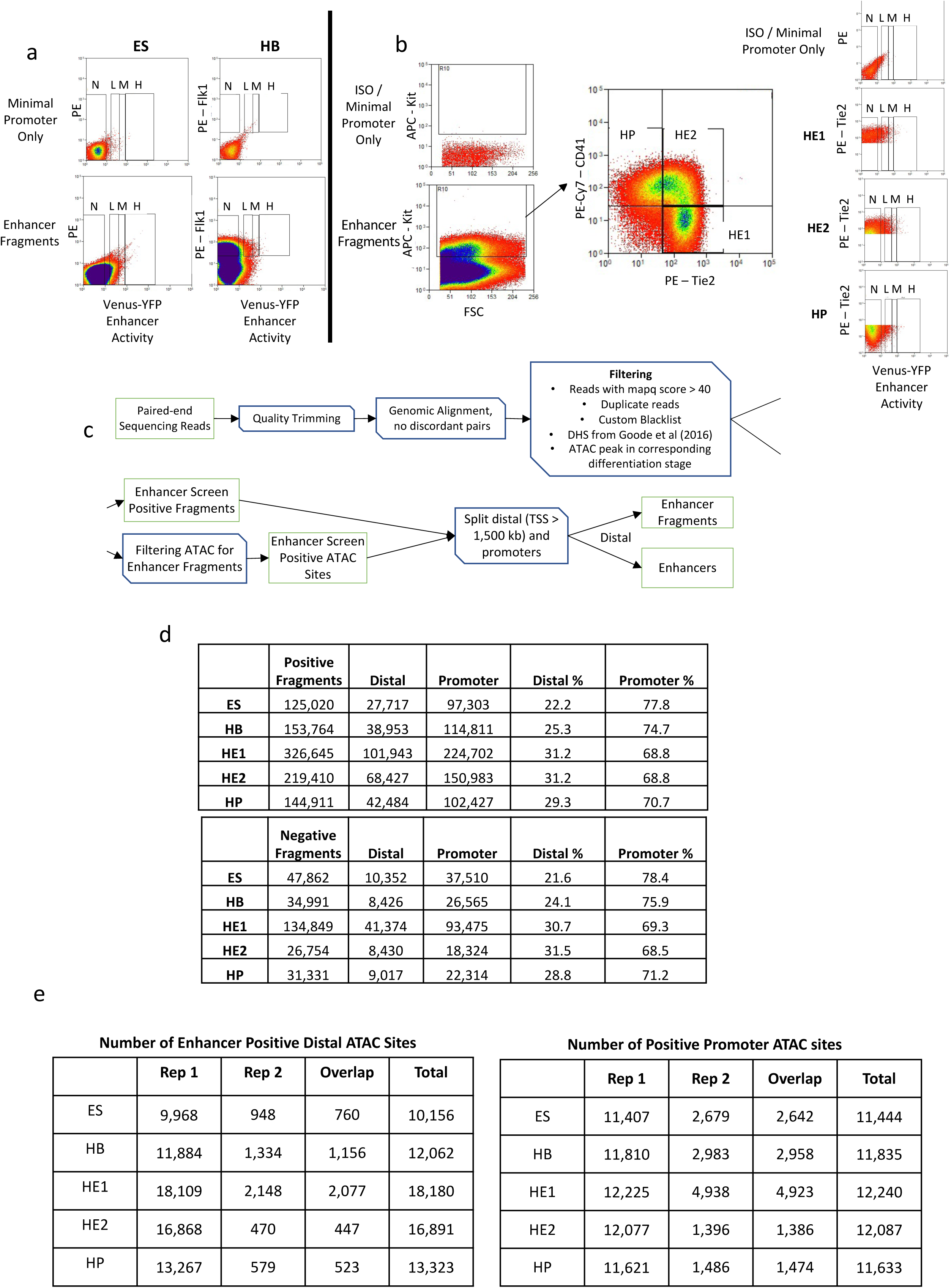
Number of stage specific enhancer and promoter fragments scoring positive in the assay. a,b) Sorting and analysis of cells with activated reporter construct, a) example of detection of enhancer activity as compared to the minimal promoter for ESCs and HB, b) representative sorting picture for all differentiation stages and gating strategy (right panel), c) Filtering pipeline, d) total number of positive and negative distal and promoter fragments recovered in the assay, e) Number of distal and promoter ATAC-sites recovered in two replicates of the assay

**Extended Figure S2:**
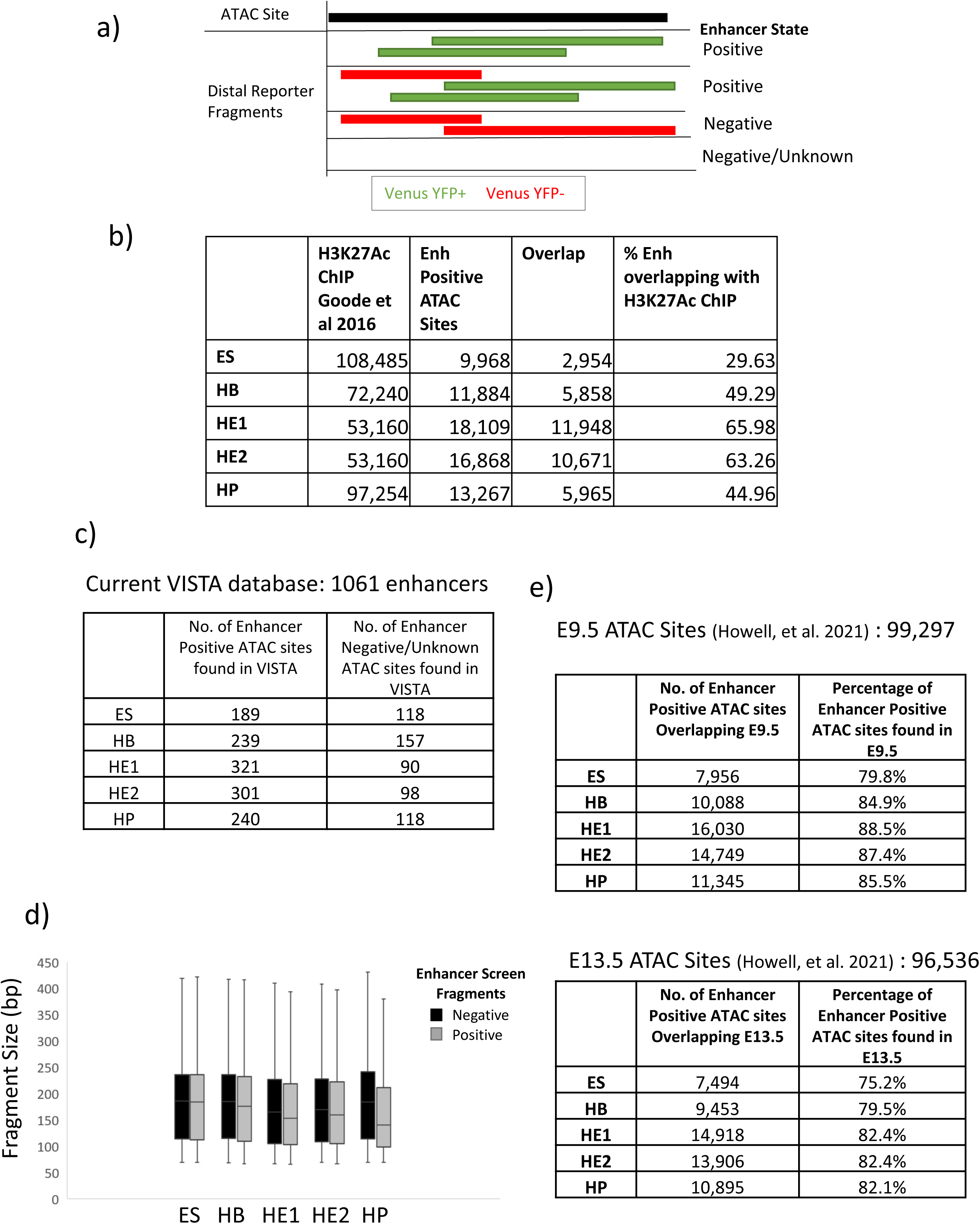
Enhancer characteristics. a) Example of a hypothetical ATAC site showing 4 possible outcomes for the specification of different types of open chromatin fragments and their correlation with the YFP signal. Fragments are labelled as negative if they do not stimulate gene expression, fragments are labelled as unknown if they are present in the original ATAC-Seq library but are neither present in the positive nor the negative fraction. b) Number of enhancers overlapping with H3K27Ac marked chromatin sites (Data from Goode at el., 2016)^22^; c) Number of enhancer positive fragments found in the VISTA database; d) size of positive and negative fragments detected in the assay (line shows median value); e) overlap of enhancer positive fragments with open chromatin sites in the hemogenic endothelium of mouse embryos (data from ^27^)

**Extended Figure S3:**
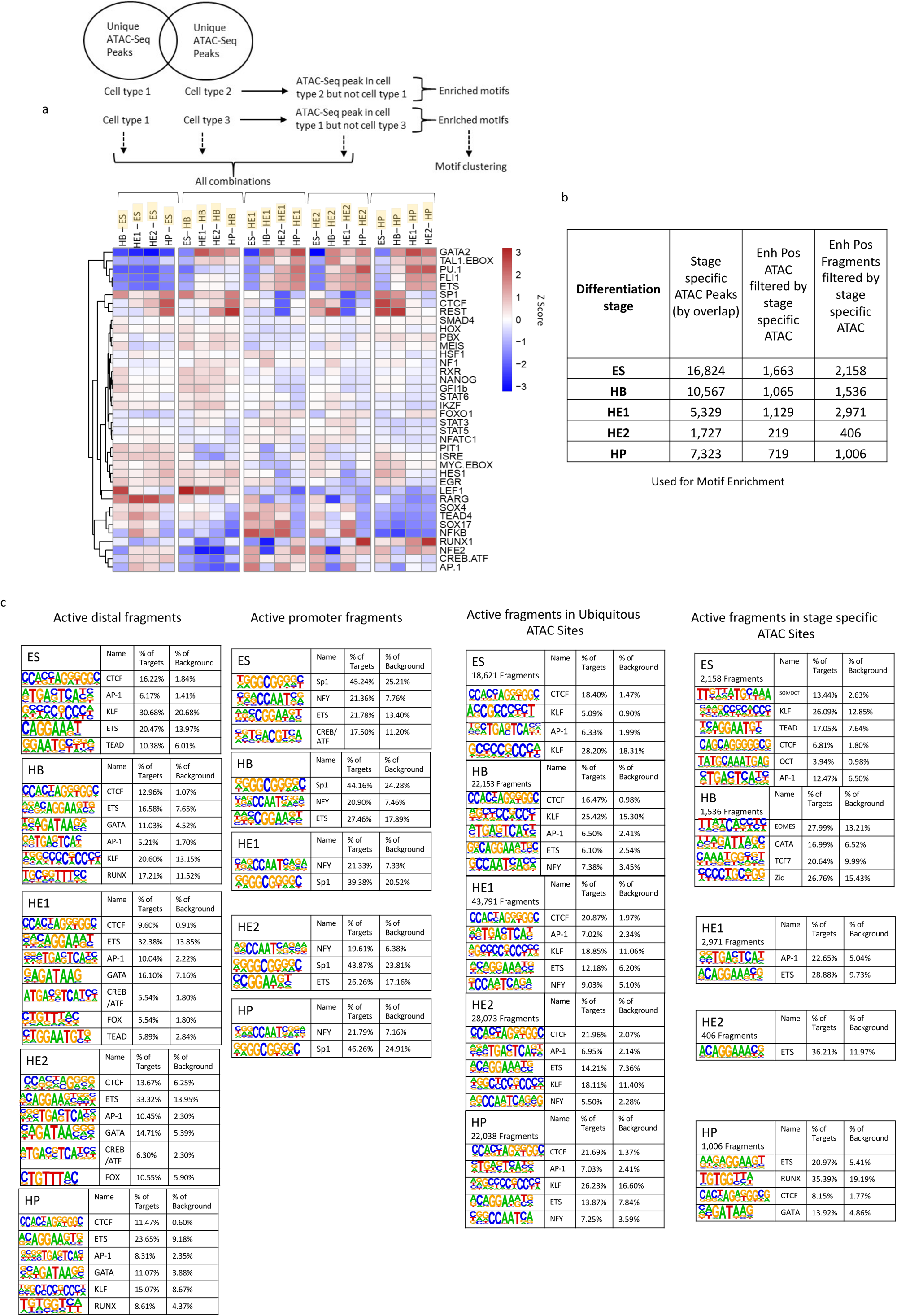
Dynamics of TF binding motif enrichment in cell type specific accessible chromatin sites, enhancer and promoter elements. (a) Identification of specific TF binding motifs in cell type specific open chromatin regions (distal elements only) as shown in ^18^. The upper panel shows the filtering strategy. The lower panel shows how specific binding motifs are enriched in ATAC-Sites that are specific for each cell type (indicated by brackets). b) Number of stage specific ATAC peaks and ATAC fragments with enhancer activity co-localizing with stage specific ATAC sites, c) TF binding motif enrichment analysis in the indicated elements using a HOMER software ^58^.

**Extended Figure S4:**
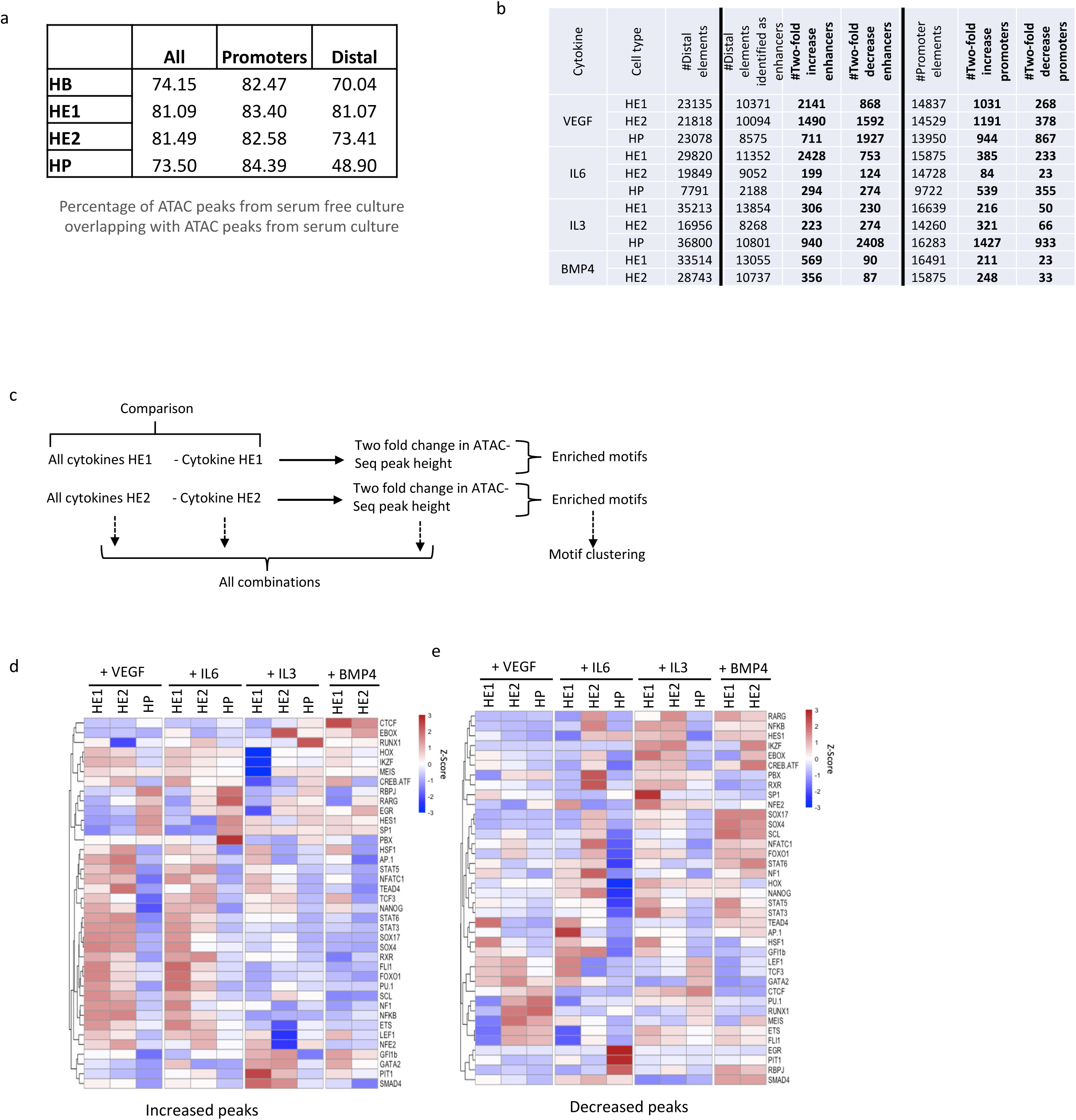
Identification of cytokine responsive enhancer elements. a) Percentage of ATAC peaks from serum free culture overlapping with ATAC peaks from serum culture; b) number of cytokine-responsive distal and promoter ATAC-peaks with enhancer activity as compared to the total number of distal and promoter elements; c,e) Motif enrichment analysis of all distal ATAC peaks irrespective of enhancer activity, c) overview of motif enrichment strategy, d-e): TF binding motif enrichment in distal ATAC peaks that are increased (d) or decreased (e) in the presence of the indicated cytokines.

**Extended Figure S5:**
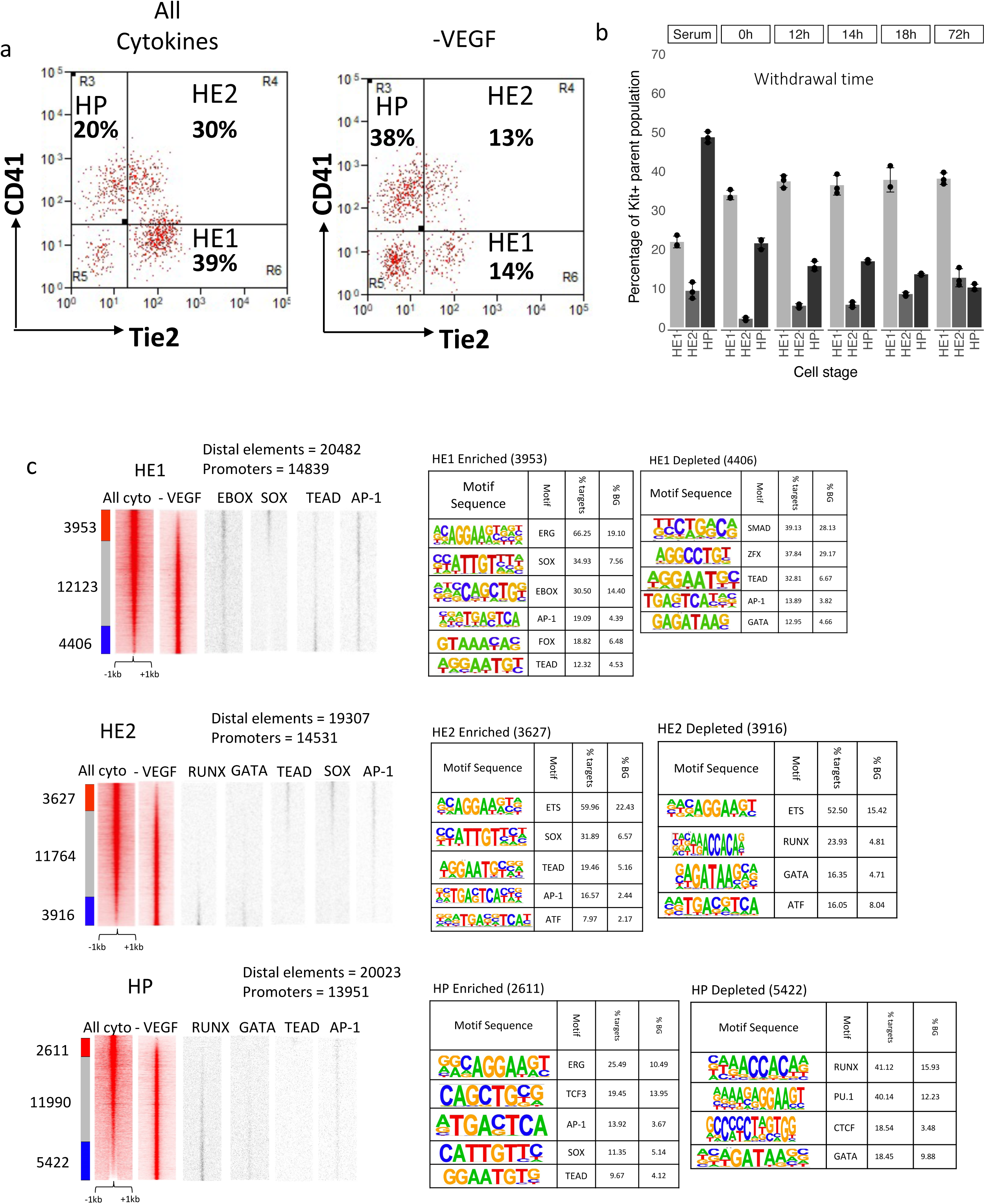
VEGF suppresses blood progenitor development. a) Representative FACS analysis of cKIT+ cells stained with antibodies against Tie2 and CD41 in the presence and absence of VEGF highlighting the percentage of HE1, HE2 and HP cells in the cultures. b) Proportion of HE1, HE2 and HP cells after VEGF was withdrawn at different times during differentiation culture from the beginning of blast culture onwards. Error bars represent the standard deviation from n=3 with the spread being indicated by dots. c) ATAC-Seq profiles of VEGF and VEGF-withdrawal cultures. The fold-difference for each peak for each condition was calculated and peaks were plotted alongside according to the fold difference to highlight ALL Cytokine and VEGF-withdrawal specific peaks. Enriched binding motifs for the indicated TFs in the peaks are plotted alongside. Right panels: Motif enrichment analysis in the specific peak populations using HOMER.

**Extended Figure S6.**
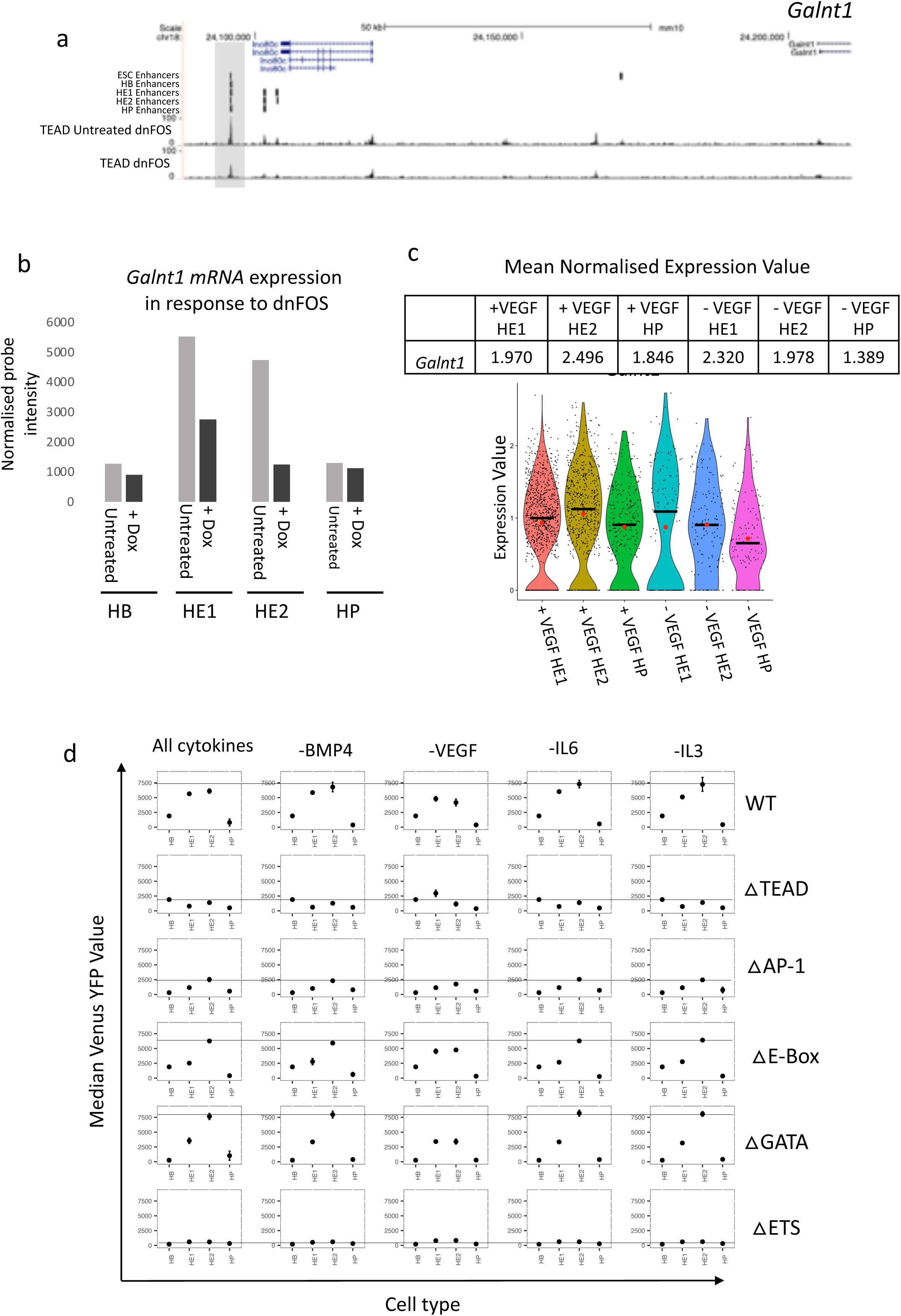
Interaction between TEAD4 and AP-1 factors at the *Galnt1* enhancer - AP-1 is required for TEAD4 binding and gene expression. a): UCSC browser screenshot showing the binding of TEAD4 to the *Galnt1* enhancer in the presence or absence of a dominant negative FOS peptide; b): *Galnt1* mRNA expression with and without dnFOS (merged duplicate RNA-Seq data from^20^). c) *Galnt1* expression in the HE1 / HE2 populations as measured by scRNA-Seq data (see below). d) Reporter gene activity driven by wild type and mutated *Galnt1* enhancer elements in the presence and absence of the indicated cytokines measured as median fluorescence of all cells. The horizontal line represents the baseline measured in the presence of all cytokines. Error bars represent the standard deviation from n=3.

**Extended Figure S7:**
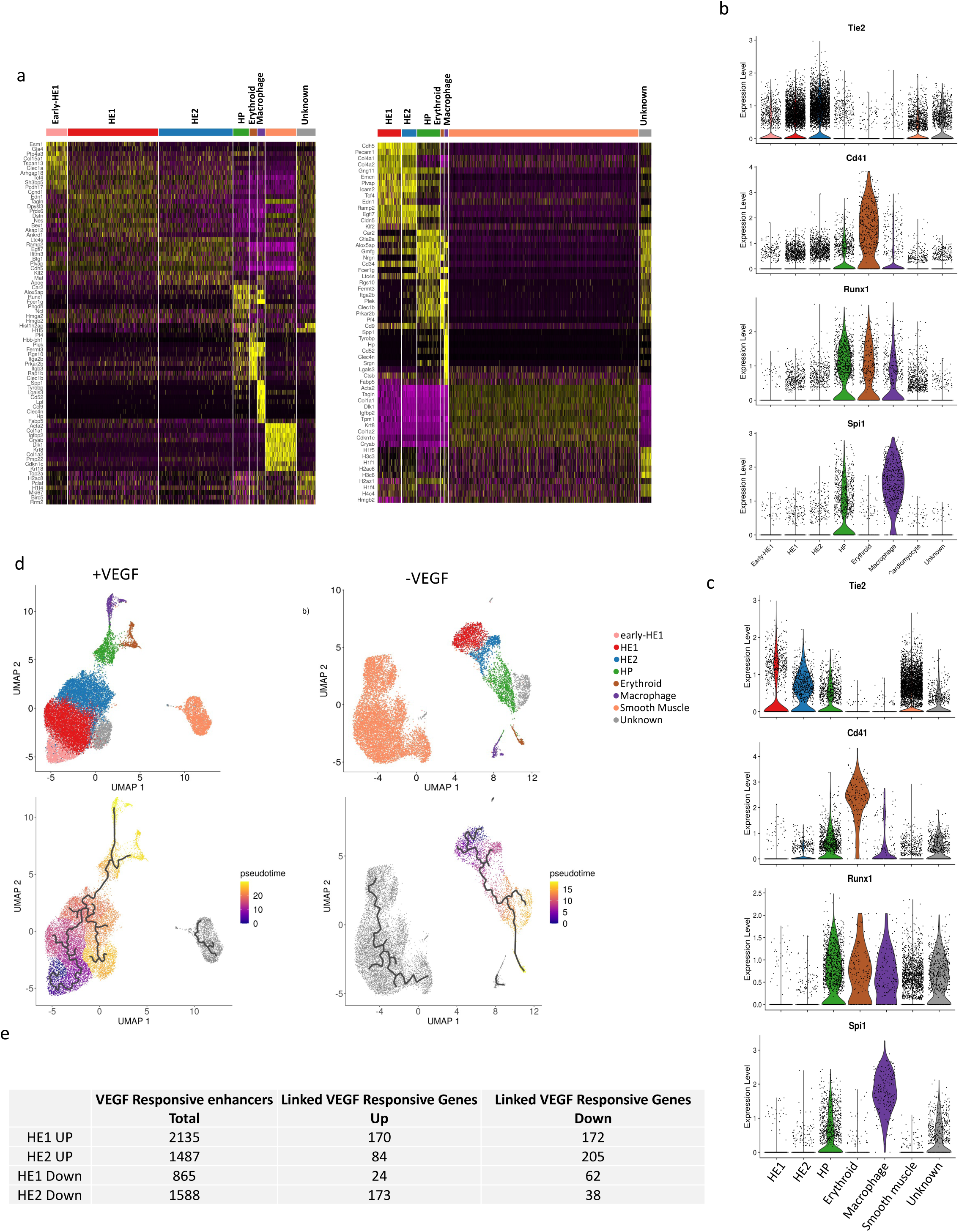
Single cell RNA-Seq analysis of hemogenic endothelium cultures with and without VEGF. Marker gene distribution in the different clusters shown in Figure 7a. b,c) Expression levels of the indicated marker genes in ES-derived immature and mature cell populations from All Cytokine (b) and VEGF withdrawal (c) cultures, demonstrating the purity of the clusters. e) Pseudotime trajectory analysis of cells derived from All Cytokine (left panel) and VEGF withdrawal cultures (right panel) of the clusters depicted in Figure 7b. f) Number of VEGF responsive genes linked to VEGF-responsive enhancer elements as measured by a 2-fold change in the ATAC-Seq signal. Expression was measured by using aggregated single cell expression values for the purified HE1 and HE2 populations using a threshold of 0.25 Log2 fold change as the currently accepted threshold for sc data.

## Methods

### ES cell culture and cell purification

The HM-1 targeting ES cell line is described in Wilkinson et al (2013)^23^. Standard ES cell differentiation is serum and the purification of differentiating cells was conducted essentially as described in Obier et al.^20^ Serum Free I.V.D Culture Embryoid bodies (EBs) were generated from HM-1 mouse embryonic stem cells by plating at 5.0×10^5^ cells/ml in serum free (SF) media into petri-grade dishes. BMP4 was added to a concentration of 5ng/ml. Cultures were left to incubate at 37°C and 5% CO_2_ for 60 hours before bFGF and Activin A were added at a concentration of 5ng/ml each. The cells were incubated for 16 hours at 37°C 5% CO2 and then sorted for FLK1+ cells as described in ^20^. FLK1+ cells were plated in serum free media on 0.1% gelatine coated plates or for larger cultures flasks at 2.25×10^4^ cells per cm. BMP4, Activin-A and bFGF were added to a concentration of 5ng/ml for 16 hours. Media was then removed, the blast culture was washed with PBS and fresh SF media was added containing BMP4 (5ng/ml), VEGF (5ng/ml), TPO (5ng/ml), SCF (100ng/ml), IL6 (10ng/ml) and IL3 (1ng/ml). For cytokine withdrawal experiments, one of BMP4, VEGF, IL6 or IL3 was not added at this stage. Blast cultures were left to incubate at 37°C and 5% CO_2_ for 72 hours before cells were harvested for cell sorting and FACS analysis. Inhibition of trypsin was achieved using Trypsin Inhibitor (Thermofisher) following the manufacturer’s instructions.

### Enhancer Reporter Library Cloning

A genome-wide enhancer reporter assay was designed based on an enhancer reporter system designed by Wilkinson et al. (2013). Briefly the enhancer reporter functions by inserting a fragment of interest upstream of a HSP68 minimal promoter and Venus-YFP reporter by Gateway^®^ cloning. The reporter construct is then transfected into the HM-1 ES cell line which has a non-functional HPRT locus. Using HPRT homology arms the report cassette becomes integrated into the HPRT locus by homologous recombination, also repairing the locus, enabling selection of clones with successful recombination. This enhancer reporter system was modified and optimised for genome-wide screening. To obtain genome-wide enhancer fragments for cloning into the reporter we isolated tn5 tagmented open-chromatin fragments based on the ATAC-Seq protocol ^33^ from cells of the five differentiation stages. Cells were obtained (2.5×10^5^) from each differentiation stage (ES, HB, HE1, HE2, HP) as detailed in the serum IVD method. Cells were pelleted in 5 aliquots of 50×10^3^ cells and following the ATAC-Seq protocol each cell pellet had a transposition mix added (2x TD Buffer 25µl, TN5 Transposase 2.5 µl, PBS 16.5 µl, 1% Digitonin 0.5 µl, 10 % Tween-20 0.5 µl, H_2_O 5 µl) and cells were gently resuspended by pipetting. The transposition reaction was then incubated in a shaker at 700 RPM, 37°C for 30 minutes. The 5 reactions were then combined and DNA purified using the Qiagen MinElute^®^ Reaction Clean-up kit and eluted in 26.5 µl H_2_O. One fifth of the purified reaction from each stage was used to produce ATAC-Seq libraries (see ATAC-Seq method) for direct sequencing and the remainder had linkers added incorporating AttB Gateway^®^ cloning sites to enable insertion into a Gateway^®^ donor vector (pDONR 221).

### Enhancer Sequencing Data Analysis

Paired end sequencing reads were trimmed using Trim Galore (Krueger F, 2021) with the parameters --nextera --length 70 –paired. The reads were then aligned to the mm10 genome using bowtie2 ^59^ using parameters --very-sensitive --fr --no-discordant -X 600 --no-mixed. Aligned reads were filtered for only those which were properly paired and with a mapq score of over 40 using Samtools and output as a bedpe file. This was further converted to a bed file of enhancer fragments by taking the first co-ordinate of read 1 and the final co-ordinate of read 2 for each pair. Duplicate fragments were removed using the uniq function and fragments were further filtered using the intersect function on Bedtools ^60^ for any which had 100% overlap with an off target background fragment list which was created by producing a library from un-transfected cells. The fragments were then filtered using Bedtools intersect for those which were found in open chromatin, defined by DNaseI-Seq, in Goode et al (2016)^18^ and further by a corresponding ATAC peak in the correct differentiation stage in HM-1 cells. Finally, the enhancer fragments were annotated using the annotatePeaks function of Homer for those within 1.5 kb of a TSS (promoter fragments) and distal fragments representing genuine enhancers. This produced a final list of enhancer fragments which was used for all further fragment analysis and by overlapping ATAC sites with enhancer fragments we defined full enhancers.

### Generation of cell lines carrying individual reporter constructs

A putative enhancer for the *Galnt1* gene containing multiple transcription factor binding sites was identified from DNaseI hypersensitive site data^18^. To test the functionality of the enhancer throughout differentiation the same enhancer reporter system ^23^ as used for the genome-wide screen was employed. The genomic sequence for the enhancer was obtained and flanking attB1 and attB2 sites were added to enable Gateway^®^ cloning. This fragment was then synthesised as an Invitrogen GeneArt Strings DNA Fragment (Thermo Fisher). The resulting fragment was cloned by the one step Gateway protocol whereby in a single reaction the fragment was interested into pDONR 221 to generate an attL-flanked entry clone by BP clonase and from these vectors into the pSKB GW-Hsp68-Venus reporter vector by LR clonase. Briefly this reaction consisted of 100 ng of Enhancer DNA, 75 ng pDONR 221, 75 ng pSKB GW-Hsp68-Venus, TE pH 8.0 to 6 µl total volume, 1.5 µl LR clonase and 0.5 µl BP clonase. The reaction was incubated at 25°C for 3 hours and then had 0.2 µg proteinase K added and a further incubation for 10 minutes at 37°C to stop the reaction. Competent DH5α bacteria (NEB) were transformed with 1 µl of Gateway^®^ reaction by incubation on ice for 30 minutes and heat shock at 42 °C for 45 seconds. The bacteria were then incubated for 1 hour at 37°C with SOC media and plated onto agar plates containing 100 µg/ml ampicillin. After overnight incubation at 37 °C colonies were picked for mini-prep cultures and mini-preps were performed using the Qiaprep^®^ Miniprep kit (Qiagen). The insert in the resulting plasmid was sanger sequenced (Source Biosciences) to check the sequence. Following confirmation of the insert sequence the same bacterial colony was used to grow cultures for Maxi-prep which was performed using the EndoFree^®^ Plasmid Maxi Kit (Qiagen).

HM-1 cells were cultured and transformed in the same way as with the Enhancer screen method. Following 6-TG treatment 5×10^6^ HM-1 cells were transfected using a Nucleofector^®^-4D (Lonza) with the P3 Primary Cell X kit. The cells were then selected for those with successful integration of the reporter cassette by treatment for 5 days with 1 x HAT containing media. At 9 to 12 days after colonies appeared clones were picked and replated on a gelatinised 96 well plate. The clones were the grown in media containing 1 x HT for 2 passages and then used for IVD and flow cytometry as detailed.

To study the impact of different transcription factor binding on the Galnt1 enhancer, sequences were produced where various transcription factor binding motifs were mutated. These mutant versions of the enhancer were cloned in the same way as with the original *Galnt1* enhancer sequence.

#### ATAC-Sequencing

ATAC-Seq was performed as described in Buenrostro et al ^61^, briefly 5000-50,000 HB, HE1, HE2 and HP cells were sorted by FACS and transposed in 1x tagment DNA buffer Tn5 transposase and 0.01% Digitonin for 30 minutes incubated at 37°C with agitation. DNA was purified using a MinElute Reaction Cleanup Kit and DNA was amplified by PCR using Nextera primers.

#### Single cell sorting and RNA-Seq

7.0×10^6^ cells from the All cytokines condition blast culture and 2.8×10^6^ cells from the VEFG withdrawal blast culture were taken for cell sorting and sc-RNA-Seq library preparation. Both samples were stained with CD41-PECY7 (1:100), KIT-APC (1:100) and TIE2-PE (1:200) and then sorted into HE1 (KIT+, TIE2+, CD41-) and HE2 (KIT+, TIE2+, CD41+) populations. From the All cytokines sample, 758000 HE1 and 380000 HE2 cells were sorted. From the VEGF withdrawal sample 71500 HE1 and 33300 HE2 cells were purified by cell sorting. Cells were resuspended in 80 μL at a concentration of 1000-1200 cells/μL for evaluation of cell viability. The viability of the All cytokines populations as measured by Trypan Blue staining was found to be 73% for HE1 and 92% for HE2. The viability of the VEGF withdrawal populations was found to be 73% and 75%. For each sample, 10,000 single cells were loaded on a Chromium Single Cell Instrument (10x Genomics) and processed.

### Data analysis methods

#### ATAC-seq Data Analysis

Single-end reads from ATAC-seq experiments were processed with Trimmomatic (version 0.39) and aligned to the mm10 mouse genome using Bowtie2 2.4.4 ^59^ using the options -very-sensitive-local. Open chromatin regions (peaks) were identified using MACS2 2.2.7.1 ^62^ using the options -B -trackline -nomodel. The peak sets were then filtered against the mm10 blacklist ^63^. Peaks were then annotated as either promoter-proximal if within 1.5kb of a transcription start site and as a distal element if not. To conduct differential chromatin accessibility analysis, a peak union was first constructed by merging peaks from two comparisons. Tag-density in a 400-bp window as centred on the peak summits was derived from bedGraph files from MACS2 peak calling using the annotatePeaks.pl function in Homer 4.11 ^58^ (with the options -size 400 -bedGraph. Tag-densities were then normalised as counts-per-million (CPM) in R v3.6.1 and further log_2_-transformed as log_2_(CPM+1). A peak which was two-fold increase or decrease different to the control was taken as being differentially accessible. A de-novo motif analysis was performed on sets of gained and lost peaks using the find -MotifsGenome.pl function in Homer using the options -size 200 -noknown. Tag density plots were constructed by retrieving the tag-density in a 2kb window centred on the peak summits with the annotatePeaks.pl function in Homer with the options -size 2000 -hist 10 -ghist -bedGraph. These were then plotted as a heat map using Java TreeView v1.1.6 ^64^.

#### Gene annotation using HiC data

Promoter capture HiC data from ESCs cells were downloaded from Novo CL et al. (accession numbers: GSM2753058, runs: SRR5972842, SRR5972842, SRR5972842, SRR5972842, SRR5972842)^34^ and from HPC7 from Comoglio et. al^65^ (accession numbers: GSM2702943 & GSM2702944, runs SRR5826938, SRR5826939). The CHi-C paired-end sequencing reads were aligned to the mouse genome mm10 build using HiCUP pipeline^66^. Initially, the raw sequencing reads were separated and then mapped against the reference genome. The aligned reads were then filtered for experimental artefacts and duplicate reads, and then re-paired. Statistically significant interactions were called using GOTHiC package^67^ and HOMER software ^58^. This protocol uses a cumulative binomial test to detect interactions between distal genomic loci that have significantly more reads than expected by chance, by using a background model of random interactions. This analysis assigns each interaction with a p-value, which represents its significance. The union of all CHiC interactions from both ESC and HPC7 cells were used to annotate positive enhancer to their related promoter.

#### Motif co-localization analysis

Genomic co-ordinates for each transcription factor (TF) binding motif were retrieved from the sets of stage specific positive enhancer fragments and from within all distal ATAC-Seq peaks using the annotatePeaks.pl function in Homer and exported as a BED file using the -mbed option. Motif co-occurrence was then measured for each stage (ES, HB, HE1, HE2 and HP) by counting the number of times a pair of TF motifs were found within 50bp of each other in the set of specific positive enhancer fragments.

To assess the significance of this co-occurrence, we carried out a re-sampling analysis whereby a number of ATAC-Seq sites equal to the number of stage specific enhancer fragments was randomly sampled from the set of all distal ATAC sites found in that stage. The number of motif pairs was then counted in this random set. This procedure was repeated 1000 times and resulted in a distribution of motif pair counts for each pair of TF binding motifs. A z-score was then calculated for each motif pair using as:

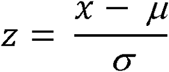

Where x is the number of motif pairs found in the specific positive enhancer fragments, μ is the average number of motif pairs in 1000 random samples, and σ is the standard deviation for those motif pair counts. A positive z-score in this case suggests that the number of motif pairs found in the positive enhancer fragments is greater than could be expected by chance. The resulting z-score matrix was then hierarchically clustered using complete linkage of the Euclidean distance in R and displayed as a heatmap.

#### Relative motif Enrichment Analysis from ATAC-Seq data

To identify transcription factor binding motifs which were enriched in a set of peaks relative to another, we calculated a relative enrichment motif score, S_ij_ for each motif i in each peak set j as: 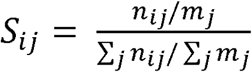, where n_ij_ is the number of instances of motif i in peak set j and m_j_ is the total number of sites in peak set j. This score was calculated for each TF motif in each of the peak sets and a matrix of enrichment scores was produced which was then hierarchically clustered using complete linkage of the Euclidean distance in R and displayed as a heat map with results scaled by either row or column.

#### Single cell RNA-Seq analysis

Fastq files from scRNA-Seq experiments were aligned to the mouse genome (mm10) using the count function in CellRanger v6.0.1 from 10x Genomics and using gene models from Ensembl (release 102) as the reference transcriptome. The resulting Unique Molecular Identifier (UMI) count matrices were processed using the Seurat package v4.0.5 ^68^ in R v4.1.2. Cell quality control was carried out for each of the four samples sequenced individually in order to remove cells which had few or a higher than expected number of detected genes or that had a high proportion of UMIs aligned to mitochondrial transcripts or quality control parameters for each sample. Genes detected in less than 3 cells were removed from the analysis.

The filtered data objects for each cell type were then combined according to their VEGF status and normalized using the LogNormalize method. In order to calculate the cell cycle stage for each of the cells, the in-built S-phase and G2M-phase marker gene lists from Seurat were first converted from human gene symbols to their corresponding mouse orthologs using the biomaRt package v2.50.0 ^69^ in R. Cell cycle stage was then inferred using the CellCycleScoring function in Seurat. The possible effect of cell cycle stage on downstream analysis was then removed from the dataset by linear regression using the ScaleData function. Clustering of cells was performed using the first 20 principal components and visualized using the Uniform Manifold Approximation and Projection (UMAP) method. Cell marker genes for each of the clusters identified were calculated using the FindAllMarkers function. A gene was considered a marker gene if it was expressed with a log2 fold-difference of 0.5 between the cluster being considered and all other cells as well as being detected in at least 50% of cells in that cluster. A gene was considered statistically significant if it had a Bonferroni adjusted p-value < 0.05. Cell type was then inferred for each cluster identified by manual inspection of the marker gene lists and comparing these to the expression known surface markers genes (Tie2, Cd41) for HE1, HE2 and HP.

Cell trajectory (pseudotime) analysis was carried out using Monocle v3.1.0 ^70^. Data from Seurat was exported to Monocle using the seuratwrappers package in R (https://github.com/satijalab/seurat-wrappers). Trajectories were then inferred using the learn graph function and ordered along pseudotime by selecting a root node which corresponded to the earliest cell-type (HE1).

## References

1. Cockerill, P.N. Structure and function of active chromatin and DNase I hypersensitive sites. FEBS J 278, 2182–210 (2011).

2. Edginton-White, B. & Bonifer, C. The transcriptional regulation of normal and malignant blood cell development. FEBS J (2021).

3. Field, A. & Adelman, K. Evaluating Enhancer Function and Transcription. Annu Rev Biochem 89, 213–234 (2020).

4. Banerji, J., Rusconi, S. & Schaffner, W. Expression of a beta-globin gene is enhanced by remote SV40 DNA sequences. Cell 27, 299–308 (1981).

5. Bonifer, C. Developmental regulation of eukaryotic gene loci: which cis-regulatory information is required? Trends Genet 16, 310–5 (2000).

6. Heintzman, N.D. et al. Distinct and predictive chromatin signatures of transcriptional promoters and enhancers in the human genome. Nat Genet 39, 311–8 (2007).

7. Rada-Iglesias, A. et al. A unique chromatin signature uncovers early developmental enhancers in humans. Nature 470, 279–83 (2011).

8. Kim, T.K. et al. Widespread transcription at neuronal activity-regulated enhancers. Nature 465, 182–7 (2010).

9. Ernst, J. & Kellis, M. Chromatin-state discovery and genome annotation with ChromHMM. Nat Protoc 12, 2478–2492 (2017).

10. Creyghton, M.P. et al. Histone H3K27ac separates active from poised enhancers and predicts developmental state. Proc Natl Acad Sci U S A 107, 21931–6 (2010).

11. Hou, T.Y. & Kraus, W.L. Spirits in the Material World: Enhancer RNAs in Transcriptional Regulation. Trends Biochem Sci 46, 138–153 (2021).

12. Dogan, N. et al. Occupancy by key transcription factors is a more accurate predictor of enhancer activity than histone modifications or chromatin accessibility. Epigenetics Chromatin 8, 16 (2015).

13. Gasperini, M. et al. A Genome-wide Framework for Mapping Gene Regulation via Cellular Genetic Screens. Cell 176, 377–390 e19 (2019).

14. Medvinsky, A.L., Samoylina, N.L., Muller, A.M. & Dzierzak, E.A. An early pre-liver intraembryonic source of CFU-S in the developing mouse. Nature 364, 64–7 (1993).

15. de Bruijn, M.F., Speck, N.A., Peeters, M.C. & Dzierzak, E. Definitive hematopoietic stem cells first develop within the major arterial regions of the mouse embryo. EMBO J 19, 2465–74 (2000).

16. Orkin, S.H. & Zon, L.I. Hematopoiesis: an evolving paradigm for stem cell biology. Cell 132, 631–44 (2008).

17. Ditadi, A., Sturgeon, C.M. & Keller, G. A view of human haematopoietic development from the Petri dish. Nat Rev Mol Cell Biol 18, 56–67 (2017).

18. Goode, D.K. et al. Dynamic Gene Regulatory Networks Drive Hematopoietic Specification and Differentiation. Dev Cell 36, 572–87 (2016).

19. Lancrin, C. et al. The haemangioblast generates haematopoietic cells through a haemogenic endothelium stage. Nature 457, 892–5 (2009).

20. Obier, N. et al. Cooperative binding of AP-1 and TEAD4 modulates the balance between vascular smooth muscle and hemogenic cell fate. Development 143, 4324–4340 (2016).

21. Vijayabaskar, M.S. et al. Identification of gene specific cis-regulatory elements during differentiation of mouse embryonic stem cells: An integrative approach using high-throughput datasets. PLoS Comput Biol 15, e1007337 (2019).

22. Huber, T.L., Kouskoff, V., Fehling, H.J., Palis, J. & Keller, G. Haemangioblast commitment is initiated in the primitive streak of the mouse embryo. Nature 432, 625–30 (2004).

23. Wilkinson, A.C. et al. Single site-specific integration targeting coupled with embryonic stem cell differentiation provides a high-throughput alternative to in vivo enhancer analyses. Biol Open 2, 1229–38 (2013).

24. Leddin, M. et al. Two distinct auto-regulatory loops operate at the PU.1 locus in B cells and myeloid cells. Blood 117, 2827–38 (2011).

25. Visel, A., Minovitsky, S., Dubchak, I. & Pennacchio, L.A. VISTA Enhancer Browser--a database of tissue-specific human enhancers. Nucleic Acids Res 35, D88–92 (2007).

26. Zhu, Q. et al. Developmental trajectory of prehematopoietic stem cell formation from endothelium. Blood 136, 845–856 (2020).

27. Howell, E.D. et al. Efficient hemogenic endothelial cell specification by RUNX1 is dependent on baseline chromatin accessibility of RUNX1-regulated TGFbeta target genes. Genes Dev 35, 1475–1489 (2021).

28. Wilson, N.K. et al. Integrated genome-scale analysis of the transcriptional regulatory landscape in a blood stem/progenitor cell model. Blood 127, e12–23 (2016).

29. Gilmour, J. et al. The Co-operation of RUNX1 with LDB1, CDK9 and BRD4 Drives Transcription Factor Complex Relocation During Haematopoietic Specification. Sci Rep 8, 10410 (2018).

30. Kellaway, S.G. et al. Different mutant RUNX1 oncoproteins program alternate haematopoietic differentiation trajectories. Life Sci Alliance 4(2021).

31. Lam, M.T., Li, W., Rosenfeld, M.G. & Glass, C.K. Enhancer RNAs and regulated transcriptional programs. Trends Biochem Sci 39, 170–82 (2014).

32. Barakat, T.S. et al. Functional Dissection of the Enhancer Repertoire in Human Embryonic Stem Cells. Cell Stem Cell 23, 276–288 e8 (2018).

33. Corces, M.R. et al. Lineage-specific and single-cell chromatin accessibility charts human hematopoiesis and leukemia evolution. Nat Genet 48, 1193–203 (2016).

34. Novo, C.L. et al. Long-Range Enhancer Interactions Are Prevalent in Mouse Embryonic Stem Cells and Are Reorganized upon Pluripotent State Transition. Cell Rep 22, 2615–2627 (2018).

35. Qi, Q. et al. Dynamic CTCF binding directly mediates interactions among cis-regulatory elements essential for hematopoiesis. Blood 137, 1327–1339 (2021).

36. Azuara, V. et al. Chromatin signatures of pluripotent cell lines. Nat Cell Biol 8, 532–8 (2006).

37. Wilson, N.K. et al. Combinatorial transcriptional control in blood stem/progenitor cells: genome-wide analysis of ten major transcriptional regulators. Cell Stem Cell 7, 532–44 (2010).

38. Pearson, S., Sroczynska, P., Lacaud, G. & Kouskoff, V. The stepwise specification of embryonic stem cells to hematopoietic fate is driven by sequential exposure to Bmp4, activin A, bFGF and VEGF. Development 135, 1525–35 (2008).

39. Shalaby, F. et al. Failure of blood-island formation and vasculogenesis in Flk-1-deficient mice. Nature 376, 62–6 (1995).

40. Lomeli, H. & Castillo-Castellanos, F. Notch signaling and the emergence of hematopoietic stem cells. Dev Dyn 249, 1302–1317 (2020).

41. Li, Y. et al. Inflammatory signaling regulates embryonic hematopoietic stem and progenitor cell production. Genes Dev 28, 2597–612 (2014).

42. Lundin, V. et al. YAP Regulates Hematopoietic Stem Cell Formation in Response to the Biomechanical Forces of Blood Flow. Dev Cell 52, 446–460 e5 (2020).

43. Lancrin, C. et al. GFI1 and GFI1B control the loss of endothelial identity of hemogenic endothelium during hematopoietic commitment. Blood 120, 314–22 (2012).

44. Nottingham, W.T. et al. Runx1-mediated hematopoietic stem-cell emergence is controlled by a Gata/Ets/SCL-regulated enhancer. Blood 110, 4188–97 (2007).

45. Lizama, C.O. et al. Repression of arterial genes in hemogenic endothelium is sufficient for haematopoietic fate acquisition. Nat Commun 6, 7739 (2015).

46. Clarke, R.L. et al. The expression of Sox17 identifies and regulates haemogenic endothelium. Nat Cell Biol 15, 502–10 (2013).

47. Lichtinger, M. et al. RUNX1 reshapes the epigenetic landscape at the onset of haematopoiesis. EMBO J 31, 4318–33 (2012).

48. Thambyrajah, R. et al. GFI1 proteins orchestrate the emergence of haematopoietic stem cells through recruitment of LSD1. Nat Cell Biol 18, 21–32 (2016).

49. Uenishi, G.I. et al. NOTCH signaling specifies arterial-type definitive hemogenic endothelium from human pluripotent stem cells. Nat Commun 9, 1828 (2018).

50. Lin, F.J., Tsai, M.J. & Tsai, S.Y. Artery and vein formation: a tug of war between different forces. EMBO Rep 8, 920–4 (2007).

51. Richard, C. et al. Endothelio-mesenchymal interaction controls runx1 expression and modulates the notch pathway to initiate aortic hematopoiesis. Dev Cell 24, 600–11 (2013).

52. Mirshekar-Syahkal, B. et al. Dlk1 is a negative regulator of emerging hematopoietic stem and progenitor cells. Haematologica 98, 163–71 (2013).

53. Voss, T.C. et al. Dynamic exchange at regulatory elements during chromatin remodeling underlies assisted loading mechanism. Cell 146, 544–54 (2011).

54. Bevington, S.L. et al. IL-2/IL-7-inducible factors pioneer the path to T cell differentiation in advance of lineage-defining factors. EMBO J 39, e105220 (2020).

55. Armesilla, A.L. et al. Vascular endothelial growth factor activates nuclear factor of activated T cells in human endothelial cells: a role for tissue factor gene expression. Mol Cell Biol 19, 2032–43 (1999).

56. Jia, J. et al. AP-1 transcription factor mediates VEGF-induced endothelial cell migration and proliferation. Microvasc Res 105, 103–8 (2016).

57. Wang, X. et al. YAP/TAZ Orchestrate VEGF Signaling during Developmental Angiogenesis. Dev Cell 42, 462–478 e7 (2017).

58. Heinz, S. et al. Simple combinations of lineage-determining transcription factors prime cis-regulatory elements required for macrophage and B cell identities. Mol Cell 38, 576–89 (2010).

59. Langmead, B. & Salzberg, S.L. Fast gapped-read alignment with Bowtie 2. Nat Methods 9, 357–9 (2012).

60. Quinlan, A.R. & Hall, I.M. BEDTools: a flexible suite of utilities for comparing genomic features. Bioinformatics 26, 841–2 (2010).

61. Buenrostro, J.D. et al. Single-cell chromatin accessibility reveals principles of regulatory variation. Nature 523, 486–90 (2015).

62. Zhang, Y. et al. Model-based analysis of ChIP-Seq (MACS). Genome Biol 9, R137 (2008).

63. Amemiya, H.M., Kundaje, A. & Boyle, A.P. The ENCODE Blacklist: Identification of Problematic Regions of the Genome. Sci Rep 9, 9354 (2019).

64. Saldanha, A.J. Java Treeview--extensible visualization of microarray data. Bioinformatics 20, 3246–8 (2004).

65. Comoglio, F. et al. Thrombopoietin signaling to chromatin elicits rapid and pervasive epigenome remodeling within poised chromatin architectures. Genome Res (2018).

66. Wingett, S. et al. HiCUP: pipeline for mapping and processing Hi-C data. F1000Res 4, 1310 (2015).

67. Mifsud, B. et al. GOTHiC, a probabilistic model to resolve complex biases and to identify real interactions in Hi-C data. PLoS One 12, e0174744 (2017).

68. Hao, Y. et al. Integrated analysis of multimodal single-cell data. Cell 184, 3573–3587 e29 (2021).

69. Durinck, S., Spellman, P.T., Birney, E. & Huber, W. Mapping identifiers for the integration of genomic datasets with the R/Bioconductor package biomaRt. Nat Protoc 4, 1184–91 (2009).

70. Trapnell, C. et al. The dynamics and regulators of cell fate decisions are revealed by pseudotemporal ordering of single cells. Nat Biotechnol 32, 381–386 (2014).

